# Microglia sculpt dopamine receptor-mediated adolescent behaviors in a social experience dependent manner

**DOI:** 10.64898/2026.09.29.755379

**Authors:** Madeline Clark, Emily Gershowitz, Ben Devlin, Roshen Amin, Julia Dziabis, Hannah Staley, Olivia Shamberger, Ja’nelle Kellam, Staci Bilbo

## Abstract

Adverse childhood experiences like social isolation (SI) contribute to long term negative health outcomes and the development of severe psychiatric disorder. Adolescence is a unique period of coordinated social and motivational system development driven by dopamine (DA) signaling. Microglia, immune cells of the brain, may play a unique role in regulating SI-induced DA system changes and associated behavioral dysfunction. Male microglia refine dopamine D1 receptors (D1rs) in the nucleus accumbens (NAc) of adolescent male rats shaping play behavior. Here we characterized the molecular developmental impacts of postweaning SI from P21-P38, with a resocialization period to P54. We tested the role of microglial engulfment of D1rs in mediating behavioral consequences of SI using neutrophil inhibitory factor (NIF) to block complement-mediated engulfment required for D1r pruning. We found sex specific impacts of SI on brain development and the DA system. In SI males, microglia engulf more NAc D1rs than group housed (GH) controls. Within GH animals, NAc D1r density is positively correlated to play and allogrooming. In SI animals, NAc D1r density is negatively correlated to biting behavior, and is not correlated to play or allogrooming. Pharmacologically inhibiting microglial engulfment of D1r via bilateral NAc NIF injection prevents observed D1r:behavior correlations in GH and SI animals. NIF treatment interacts with SI, exaggerating aggressive-like behavior, showing a role for microglia in regulating aggressive-like behavior via complement pruning in adolescence. Microglia may play a unique role in regulating behavior that drive negative health outcomes in the face of developmental stress, including SI.

## 1. Introduction

Adverse childhood experiences including social isolation (SI) cumulatively increase risk for serious mental health disorders, chronic illness and early mortality^1–4^. A key contributor to this early mortality is a lack of social support and connection^5^. A key feature of serious mental health disorders including schizophrenia and bipolar disorder is social difficulty^6,7^. Adverse experience-induced social dysfunction may exacerbate lack of connection and social support, reinforcing negative health outcomes^8^. During the COVID19 pandemic, 26 million adolescents aged 12-17 were impacted by a period of SI, the consequences for which we do not yet fully understand^9^.

Adolescence is a unique period of increased social, reward, and novelty seeking^10^. This is observed through increased behavioral complexity, including play fighting behaviors which are developmentally regulated in adolescence across species including rats and humans^11–15^. Play is an innately rewarding, dynamic whole body process, requiring moment to moment flexible appraisal and response^13,16,17^. Play requires cooperation in role reversal and restraint to allow reciprocal engagement; a lack of restraint and reciprocity can escalate into aggression^18^. In rats, adolescent social behavior is sexually dimorphic where males play more than females and females allogroom more than males^15^. These behaviors have lasting consequences, where adolescent females who reciprocally allogroom during a mild stressor in adolescence have decreased mammary tumors and increased overall lifespan^19^. Adolescent play fighting behavior primes resiliency in the face of threat, motor coordination, and in males allows for later reproductive success^20–22^.

Social and motivated behaviors in the brain are coordinated by the dopamine (DA) system^23,24^. During adolescence, DA rich brain regions undergo synaptic remodeling, patterning the brain for behavioral stabilization into adulthood. In humans this process can extend into the late twenties and early thirties^25–33^. Adolescence has been called a critical window for social development due to heightened plasticity and protracted development^34^. This protracted development makes this system particularly sensitive to environmental perturbations, disrupting development and contributing to psychiatric disorder. The nucleus accumbens (NAc) is a DA rich brain region of the ventral striatum involved in social behavioral expression^23^. The NAc receives DAergic input from neuronal tyrosine hydroxylase (TH) fibers of the ventral tegmental area (VTA)^23^. VTA inputs to the NAc are tonically active, with changes in DA output and firing patterns corresponding to aspects of salience including reward prediction, aversion, vigor, belief state and expectation^35–40^.

The resident macrophages of the brain, microglia, have been shown to sense and respond to DA activity in the striatum^41^. In other brain regions like the lateral geniculate nucleus, microglia use complement mediated pruning to regulate synaptic development in an activity dependent manner^42^. The complement receptor complex C3r/Cd11b on microglia recognizes C3 tagged proteins for engulfment and degradation^43^. Our group has previously found that microglia in the NAc of male rats use complement-mediated pruning to refine dopamine D1 receptors (D1rs), orchestrating a developmental decrease in receptor density from mid adolescence at postnatal days 30-38 (P30-P38) to late adolescence/early adulthood at P54^44^. This set of experiments prevented microglia complement-mediated engulfment of NAc D1rs with local injection of neutrophil inhibitory factor (NIF) which binds to and blocks CD11b^44^. NIF treatment increased D1r density and play behavior in socialized males demonstrating a microglial role in DA development and its regulation of social behaviors^44^. Microglia are remarkably sensitive to environmental perturbations; for instance, social stress induces microglial over-pruning of synapses in the rodent mPFC and release of proinflammatory cytokines, altering social approach in response to chronic social defeat^45,46^. Collectively these data suggest microglia are uniquely poised to regulate SI-induced behavioral dysfunction via NAc D1r developmental wiring in males.

SI in rats increases play and aggressive like behaviors^47–49^. SI also increases DA release from the VTA, and TH fiber density in adult animals persistently isolated from weaning^50–53^. Adolescent-specific SI has been shown to increase vigor and perseverative behaviors, preventing behavioral flexibility into adulthood^54–56^. Notably, postweaning SI has been associated with increased drug seeking for cocaine, methamphetamine and ethanol into adulthood^52,57–59^. Understanding the mechanisms of SI-induced DA dysfunction may yield novel insights into vulnerabilities that may increase addiction liability. Here we aim to outline the role of microglia in mediating adolescent SI-induced behavioral changes.

We characterize DA system development in an adolescent-specific model of SI. We recapitulate increases in play behaviors including pouncing, boxing, pinning and chasing following SI. We also see increases in aggressive-like behaviors including biting and non-reciprocal vigor, which we term “assertive play”. There are sex-specific changes in behavior and DA system development, with SI-induced shifts in NAc TH and D1r density in males and only TH density in females. Given developmental complement pruning of NAc D1rs is male specific, males were used for the majority of this study with females included for as many endpoints as feasible to further understanding. We show in males that SI increases D1r pruning by NAc microglia. This pruning by microglia mediates associations between receptor density and behavioral output in a social environment dependent manner. Play is positively correlated with NAc D1r density in group housed (GH) animals, and biting behaviors are negatively correlated with NAc D1r. Preventing microglia pruning via NIF administration disrupts the typical NAc D1r:behavior correlation and exacerbates non-reciprocal aggressive like behavior in SI.

## 2. Methods

### 2.1 General Methods

#### 2.1.1 Animals

TH:Cre rats HsdSage:SD-THem1(IRES-Cre)Sage were obtained from Envigo now Inotiv. Animals were group housed in individually ventilated cages with ad libitum access to (Purina lab Diet 5001) and water. The colony had a 12:12h light cycle with lights on at 02:00h and was maintained at 22C. Cages were changed every two weeks with spot changes as necessary. Breeder males were removed from pairs before parturition. Offspring were weaned at P21 and given supplemental care including a pellet food slurry and gel pack for hydration. GH animals were sex matched in pairs, while isolated animals were single housed with an added bone for enrichment. SI animals for experiments past P38 were rehoused together at P38. This reporting follows the ARRIVE guidelines and the National Institute of Health’s Guide for the Care and Use of Laboratory Animals. Experiments were approved by Duke University Institutional Animal Care and Use Committee.

#### 2.1.2 Play Behavior

GH experimental animals were single housed at P37, a day before testing. Conspecifics were age and sex matched, housed in groups (2-3 animals/cage). Conspecifics were littermates and never a cage mate post weaning at P21. On the day of testing, animals were habituated for 30 minutes, starting 15 minutes before lights out. Animals play by red light, where a conspecific is added to the experimental animal’s cage and their interactions recorded for 15 minutes using an infrared webcam and OBS Studio. Videos were coded and behavioral scoring was blinded. Behavior scoring was manually recorded using Solomon Coder for these behaviors: General Exploring, Bite, Bouncy Gate, Chasing, Anogenital Sniffing, Pouncing, Pinning, Boxing, Crawling Over/Under, General Social Interaction, Grooming, Assertive play and Allogrooming. Please see supplementary Video 1 for example of reciprocal play behavior and Video 2 for assertive play behavior examples.

#### 2.1.3 Social Preference and Novelty Test

Animals were single housed and habituated to the testing room and plexiglass social chamber arena on P53. Animals were kept in the testing room for 30 minutes, before being allowed to freely roam the empty social chambers for 10 minutes and then returning to the home cage. Habituation and testing on test day occurs between 08:00h and 13:30h. On testing day, P54, animals were habituated to the room for 30 minutes. Animals then undergo the social preference task where they were placed at the center of the three-chamber arena, in which a social conspecific or a dog toy is placed at either end of the apparatus. Conspecifics and toys were kept behind plexiglass bars, so that animals may sniff and interact but not freely. Experimental animals may freely roam for 5 minutes, after which time the toy is replaced with a new conspecific. Experimental animals were recentered in the arena at the beginning of the social novelty portion of the task for a subsequent 5 minutes.

#### 2.1.4 Tissue Collection and Sectioning

Animals were euthanized via CO2 ventilation and transcardially perfused with ice cold 0.9% isotonic saline followed by 4% paraformaldehyde (PFA) for fixation. Brains were post fixed for 24 hours in 4% PFA before being placed in 30% sucrose in phosphate buffered saline (PBS) with sodium azide (0.1%) until sunk, at least 3 days. Brains were frozen in 2-methylbutane on dry ice and stored at -80C.

#### 2.1.5 Immunohistochemistry

Upon sectioning, brains were brought from -80C to -25C in a cryostat chamber for 30 minutes before being serially sectioned into 40µm sections. Sections were stored in Eppendorf tubes containing cryoprotectant with sodium azide at -20C until staining. For staining, sections underwent 4×5minute 1xPBS (pH7.4) washes, blocked in 1xPBS with 10% normal goat serum, 0.3% Triton-X and 1% H2O2 for one hour. Sections incubated in a primary solution of PBS and 2% normal serum, in PBS with 2% normal serum and 0.3% Triton-X overnight at room temp. Primaries were used as follows: TH (1:1000, Immunostar #22941), D1r (1:1000 Novus NB110), IBA1 (1:1000, Wako #019-19741v, Synaptic systems #234-009), CD68 (1:200, Biolegend #137004). Sections were then rinsed in PBS 4×5minutes, and incubated with a secondary antibody either 1:500 for 4 hours or 1:250 for two hours. Sections were then rinsed in PBS 3×10minutes and then mounted onto gelatin subbed slides within 5 days. Sections were cover slipped using Vectashield plus DAPI (Vector Labs) or Fluoromount-G with DAPI (ThermoFisher).

### 2.2 Image Analysis

#### 2.2.1 Mean Grey Value

Mean grey value measurements are gross total measurements of relative protein density via fluorescent signal by proteins bound to epitope specific fluorescent antibodies. Z stack images were obtained using either Zeiss AxioImagerM2 with apotome or Fluoview V3000 confocal microscope. Images were taken using a 40x objective, or 20x with 2x optical zoom. Z-stacks were 10x.5 µm step images. Z-stacks were transformed using ImageJ open source software with preinstalled plugins (FIJI), into Max-Intensity projection 8-bit tiffs. Images were then imported as a sequence, and a background measurement was taken. Images were individually thresholded to protein signal area, and a measurement was taken of all the pixels in a thresholded area, with background subtracted from the signal measurement.

#### 2.2.2 Sholl Analysis

For Microglia 2D Sholl Analysis, at least 5 IBA1+ microglia were selected for analysis per animal across at least 2-3 images. Those individual microglia were put through the Ilastik pixel segmentation pipeline to generate binary segmentations of the images. These binary segmentations were processed using a custom python script based on the sholl analysis Fiji plugin that skeletonizes the image and plots concentric rings from the soma of each cell, measuring the intersections of those rings with cell branches as a measure of complexity.

#### 2.2.3 IMARIS reconstructions

For volumetric reconstructions for microglial engulfment of D1r, 60x Z-stack images were taken at .5µm step size through the full depth of the tissue using an Olympus FV3000 Confocal. At least 4 Z-stacks across two sections per animal were quantified. Using IMARIS v9.10, a surface was rendered using the IBA1 channel. Lysosomal marker CD68 was masked inside of the IBA1 channel. Then a surface of CD68 was generated, and a mask of D1r inside of CD68 was generated. Finally a surface was generated from D1r signal inside of the CD68+ microglia lysosomes.

For FosB quantification, similar steps as above were taken. Using an Olympus FluoView 3000 confocal, a 10 step Z stack at .5µm was taken of tissue sections stained for NeuN, FosB and IBA1. At least 4 images per animal were quantified. A mask of NeuN+ or IBA+ positive stain was created. Then FosB signal within those stains was quantified, and a watershed function was used to eliminate duplicate nuclei in the instance of cellular crowding.

#### 2.1.6 Microglia Isolation

Microglia were extracted using methods developed in our lab, for ref. see our STAR protocol^60^. In brief, animals were transcardially perfused with ice cold saline. Brains were rapidly dissected on ice, where the NAc/ventral striatum was macro dissected. Tissue was homogenized using a razor blade and placed into an enzyme digestion mix on ice until further processing. Samples were then placed into a 37C water bath, followed by mechanical dissociation. Cells were then incubated with rat CD11b conjugated magnetic beads (Miltenyi cat#130-105-643). CD11b+ microglia were selected via antibody binding and magnetic dissociation. Samples were spun down and stored in 500µl trizol in a -80C until processing.

#### 2.1.7 RNA Extraction and Microglia Sequencing

Rats were saline-perfused, and their brains dissected and split at the midline using a razor blade. The NAc/Ventral striatum was dissected, and microglia were isolated from using a CD11b antibody-based procedure, according to our protocol^60^. RNA was from CD11b+ isolated microglia using TRIzol based chloroform extraction and samples were transferred to MedGenome (Foster City, California) for library preparation and sequencing. Raw fastQ files were aligned to Rattus Norvegicus GRCr8 using STAR (v2.7.5c) and featurecounts (v1.6.3) on the Duke Compute Cluster with custom bash scripts. Genes were filtered, removing mitochondrial and ribosomal genes. Genes remaining after filtering were only included in analysis if they were present at 10 counts in at least 4 samples. Differentially expressed genes were calculated using DESeq2 in R 4.4.0. Significant genes were plotted using R and Python. Genes were run through GOrilla gene ontology analysis; however, there were not significant findings to report (data not shown).

### 2.3 Surgery

#### 2.3.1 NIF

For all surgeries, animals were weighed before being anesthetized under 4% isoflurane and brought up to 2.5-3% throughout the surgery. The rats’ heads were fitted into a Stoelting stereotaxic rig and given 5mg/kg ketofen (Zoetis) i.p. at the beginning of the surgery. Heads were shaved before betadine and ethanol preparation of the skin prior to incision. Hamilton 1µl #7001 syringes with metal needle tip (Hamilton, 80100) were used for all stereotaxic injections. P30 animals were bilaterally injected with NIF (1x reconstitution, 200µg/mL R&D systems #5845-NF-050) or sterile PBS at a rate of 50nL/min for a total of 6 minutes. Needle tips were left for one minute after injection before removing. NAc coordinates used were AP 2.3mm, ML 2.5mm, DV-6mm with a 10 degree angle for each stereotaxic arm.

#### 2.3.2 DREADD surgeries and experiment

For chemogenetic manipulation of the DA system, we expressed Cre-dependent Designer Receptors Exclusively Activated by Designer Drugs (DREADDs) in TH+ neurons using adeno-associated viruses (AAVs). P21 males received bilateral 0.5µl injection of either an excitatory (Gq), inhibitory (Gi) or mCherry (pAAV8-hSyn-hM3D(Gq)-mCherry, pAAV8-hSyn-hM4D(Gi)-mCherry, pAAV8-hSyn-mCherry, addgene) control virus into the VTA. The coordinates used were AP 4.7mm, ML 1.75mm and DV-8mm at a 10 degree angle. Animals were given 1mg/kg clozapine-N-oxide (CNO, Tocris) dissolved in 1% DMSO, daily from P27-P31. Animals were weighed and injected daily between 09:00h and 12:00h. Animals were collected 90 minutes after the final injection on P31. Viral expression was validated with imaging using an Olympus VS200 ASW 3.2.1 slide scanner at 20x.

### 2.4 Statistical Analysis

The statistics presented here were run on GraphPad Prism v10 unless otherwise specified. Animals/measures that are more than two standard deviations or more separated from the group mean were considered outliers and removed from group comparisons. Animals who had 3 or more outliers across play behaviors were removed overall (no more than 1-2 one animal per group, per experiment). Using Levine’s test for normality, many comparisons fail a normal distribution. To account for this Welch’s T-tests were performed with bonferoni-dunn multiple post hoc test. Principal component analyses were run using Prism v10, with the first 3 PC components run with Welch’s T-test and bonferonni-dunn post hoc tests. All correlations used pearson’s correlation coefficient with regression lines present for graphical representation. Two-way ANOVA and Tukey’s post hoc test was used to test for locomotor differences in DREADD groups in response to cocaine. One-way ANOVAs were run on single animal averages of protein fluorescent intensity to test for mean differences in various dopamine and microglia endpoints in DREADD manipulated brains.

Immunohistochemical data proved to follow a normal distribution. Therefore, standard two-way ANOVA tests with Tukey’s post hoc test was run for immunohistochemical data looking at factors of age and condition impacting D1r, TH and IBA1 markers across brain regions. Interaction effects were tested for if P=<.1. For FosB cell population data, all groups were not normally distributed as found through Shapiro-Wilk (Microglia cells, lower overall N) or D’agostino-pearson (Neurons, much higher N) tests using R v3.6.0. To interrogate population level differences in FosB intensity expression, a Kolmogorov-Smirnov test was used to interrogate significantly different population distributions and population median.

## 3. Results

### 3.1 Adolescent SI alters play behavioral repertoire and later social novelty seeking in a sex-specific manner

In our paradigm of adolescent SI, animals were weaned at P21 and isolated until P38, after which they were resocialized in pairs until P54 (Fig.1A). GH animals were weaned into sex matched pairs for the duration of the study. Animals undergo recorded play behavior at P38 with an age and sex matched conspecific, where the conspecific is placed into the experimental animal’s home cage for 15 minutes. Following play, animals were returned to their home cages, or resocialized, until social novelty and preference testing at P54 (Fig.1A). Play behaviors were hand scored and included pouncing, boxing, pinning, chasing, assertive play and bite (Fig.1B). Non-play social behaviors that were scored included anogenital sniffing, allogrooming, crawling over/under, and generic social investigation. Other behaviors collected included bouncy gait, grooming self and general exploratory behavior. There is a rich literature investigating facets of rat behavior that has informed selecting these behaviors^17,18,21,44^. To illustrate the complexity of behavior repertoire use, average relative percent behavior use per sex and social condition are represented in a pie chart (Fig.1C). To further understand changes in behavioral repertoire, the 15 behavioral components were analyzed by principal component analysis (PCA) (SFig.1A). Represented are animals across groups in 3D PCA behavior space showing a clear segregation of behavioral repertoire between SI and GH rats in both males and females, with greater behavioral variance overall in SI rats (Fig.1D). SI males take up 18x greater behavioral complexity space than GH males (Fig.1D), and SI females take up 13x more than GH females (Fig.1D’). Quantification of the PC values showed a significant difference of PC1 in males and females (SFig.1B,E).

**Figure 1:**
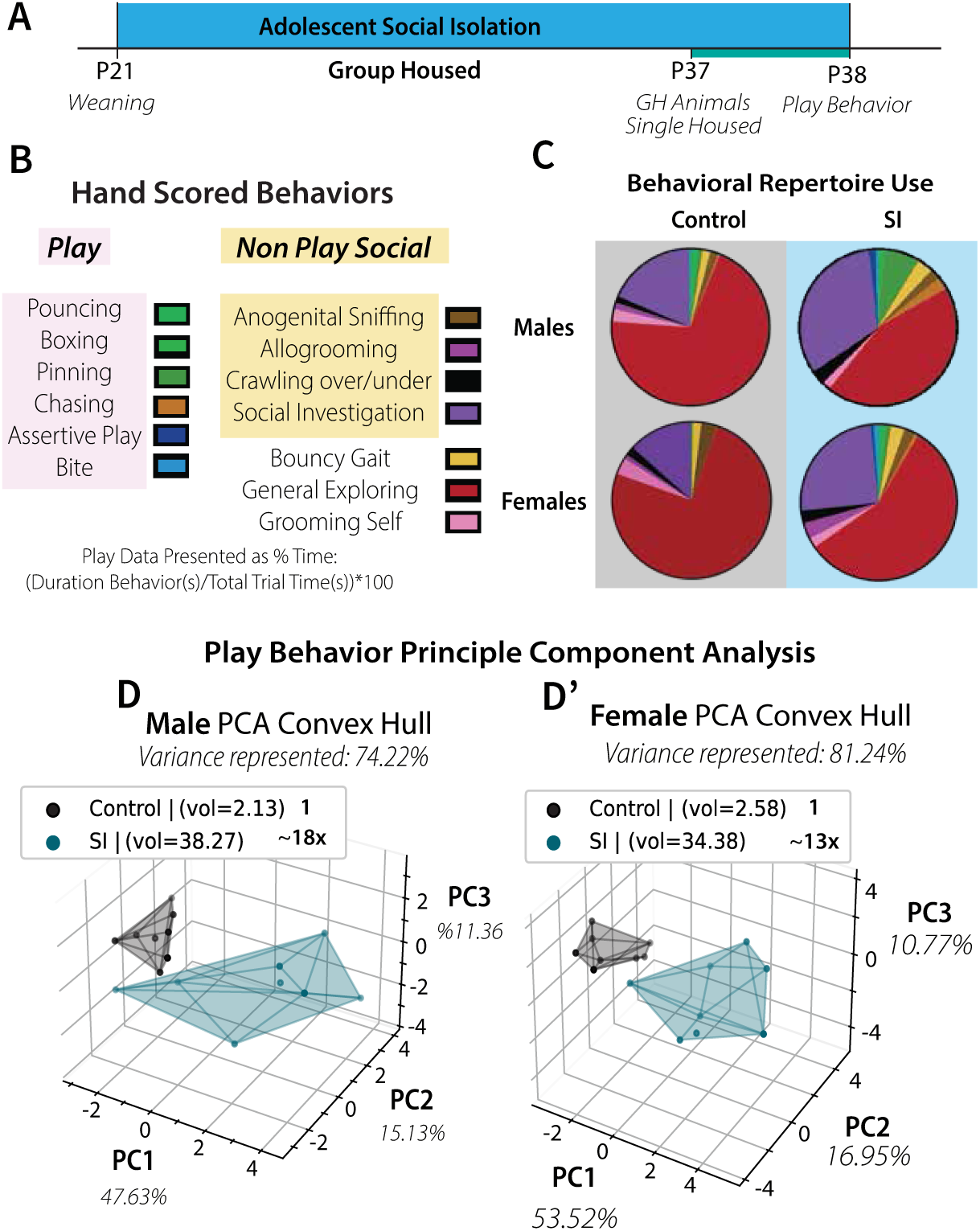
Adolescent SI Increases Behavioral Complexity in Males and Females, Persistently Increases Social Novelty in Males. A Experimental timeline of Adolescent SI, where isolated animals are singly housed at P21 until P38, and are resocialized until P54. Group housed animals are single housed 24 hours before play behavior at P38. All social partners, play, preference and novelty, are age and sex matched. B List of scored behaviors for each 15 minute play bout. C Pie charts showing the total averaged behavioral repertoire use across sex and social condition. D PCA analysis of animal behavior represented in 3D space, capturing 74.22% of the variance in males and 81.42% of the variance in females. N=8-10 animals per group.

We next analyzed individual behavior classifications. SI males and females show a significant increase in play behavior (Fig.2B). SI females and not males show a significant increase in non-play social interactions (Fig.2C). One behavior that drives this difference is allogrooming, where SI females show a significant increase that males do not (Fig.2D). Assertive play behavior is non-reciprocal behavior, violating the rule that play remains play with balanced reciprocity^18^. Biting behavior can be both playful and aggressive^15,18^. To determine the aggressive nature of biting behavior in these animals we performed a Pearson’s correlation (PCC) of % bite time and % assertive play time in males and females (Fig.2E). There is a significant correlation in males and females (Fig.2E). These two behaviors have been compiled into one aggression measure, where there is a significant increase in SI males and females (Fig.2F). Interestingly when bite behavior is normalized to play behavior, there is a significant increase in SI females (SFig.2D’). When anogenital sniffing behavior is normalized to non-play social behavior, SI animals of both sexes show decreases suggesting changes in salient social stimuli outside of play and aggression (SFig.2G’).

**Figure 2:**
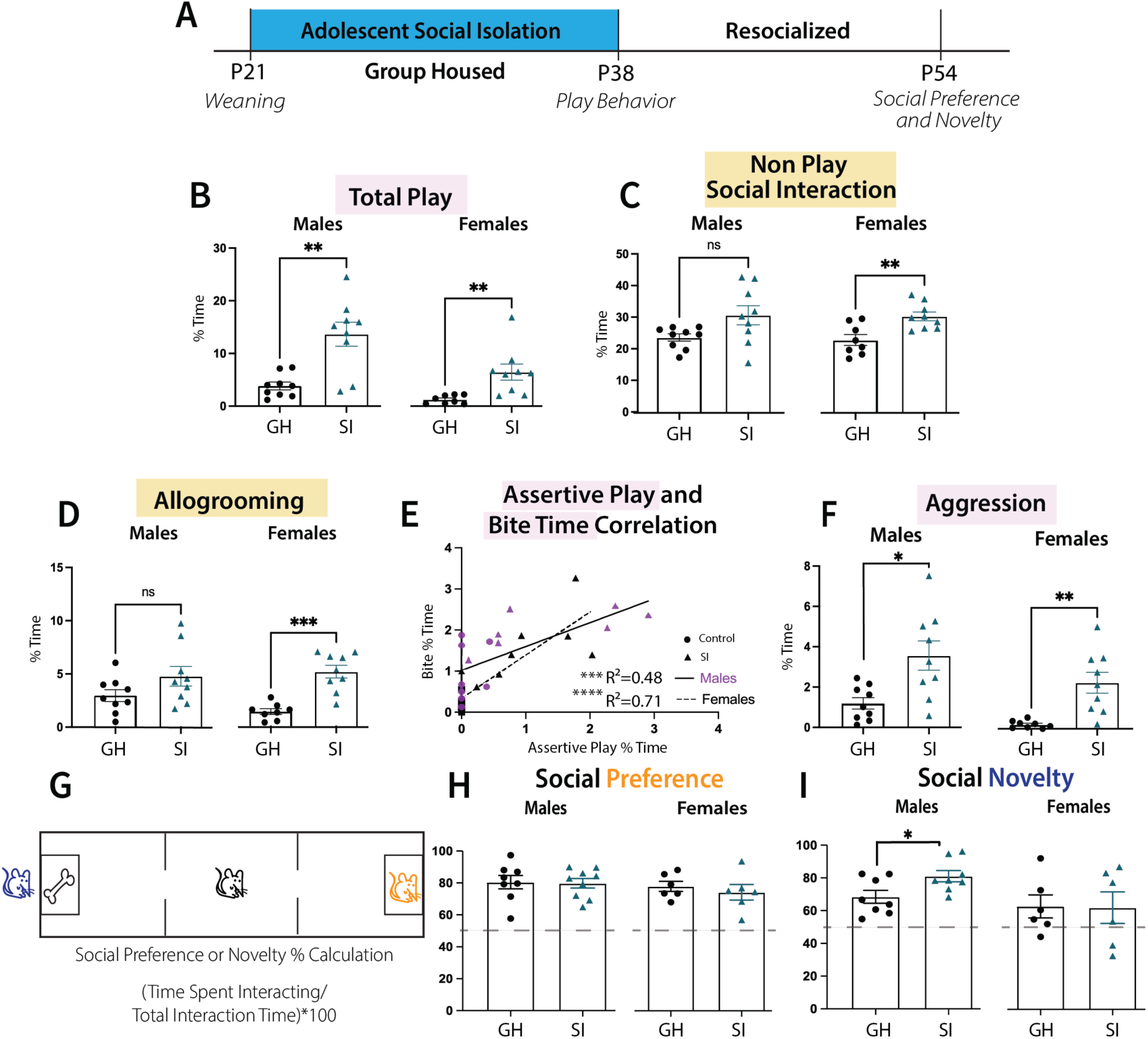
Adolescent SI Increases play, Persistently Increases Social Novelty in Males. A Experimental timeline of Adolescent SI, where isolated animals are singly housed at P21 until P38, and are resocialized until P54. B Quantification of total play behavior represented as % time of total 15 minute play bout. C Quantification of non-play social interaction. D Quantification of allogrooming behavior. E PCC of assertive play and bite time across males and females with regression lines plotted for each sex and R^2^ values presented. F Quantification of aggression, comprising assertive play and bite time. G Social preference and novelty testing where animals may freely interact with a social conspecific or a toy. After a period of five minutes the toy is replaced with a novel age and sex matched conspecific, for another period of five minutes comprising the novelty portion of the task. H Quantification of social preference and I Social Novelty. Dots represent single animals with the same animals represented across the plots shown here. *P<.05 **<.01***.001 by multiple unpaired Welch’s T-Test with Bonferroni-Dunn post hoc testing. Welch’s T-Test controls for unequal population standard deviations, while Bonferroni-Dunn post hoc testing controls for multiple tests preventing false positive errors. N=8-10 animals per group. All error bars = SEM, animals per age and condition are at least 4 distinct litters.

To understand the lasting impact of adolescent SI on social behavior animals underwent social preference and novelty testing at P54 (Fig.2A). Animals were placed in the center of a 3 chamber arena where they may freely roam to interact with a social conspecific confined to one end, or a dog toy confined to the other end for five minutes (Fig.2G). Following this time, the dog toy is replaced with a new age and sex matched conspecific for another five minutes comprising the novelty portion of the trial (Fig.2G). We observed no difference in social preference in males or females (Fig.2H). We did see an increase in social novelty specifically in males (Fig.2I). These data demonstrate persistent social changes in males following SI, while SI itself increases behavioral complexity, engagement in play, and aggressive behaviors across sexes. We next sought to understand the molecular underpinnings of these social behavioral changes through characterization of the DA system development in the NAc, a key brain region in play behavior expression^44,61^.

### 3.2 Adolescent SI alters mesolimbic DA system development in a sex specific manner

To characterize NAc DA system development in our hands, we collected brains at three developmental timepoints, P30, P38 and P54 (Fig.3A). These timepoints capture the middle (P30) and end (P38) of the isolation period, as well as a post-resocialization timepoint (P54). These timepoints also reflect key timing for microglia pruning of D1rs at P30^44^, and timepoints of previously shown play (P38) and social novelty behaviors (P54). We hypothesize there may be dynamic interactions of neuronal activity, microglia and receptor density in the NAc particularly in adolescence which we have represented here (Fig.3B). Microglia are known to sense and respond to activity of neurons regulating engulfment^62,63^. Microglia prune D1rs of the NAc, which primarily reside on local spiny projection neurons^64^. VTA DA terminals project to the NAc and are labelled by TH, the rate limiting enzyme for DA synthesis^65^. In lieu of a direct measure of activity, we quantified FosB, an activity dependent protein, in neurons and microglia^66,67^. We used an antibody (Cell Signaling cat# 2263) that also labels delta-FosB, a long lived isoform that accumulates in response to repeated activity^68^, though the isoforms are indistinguishable here. We sought to interrogate how SI alters DA system development, including shifts in activity. In males there is a significant interaction of age and SI on TH fiber input to the NAc, with an SI-induced increase at P30 that is normalized at P38 and P54 (Fig.3C). There is also a significant interaction in females, where there are trending (p=0.06) SI increases in TH fiber density at P30 and P54 (Fig.3D). There is significantly increased TH fiber density in P30 GH females compared to P38 and P54, while in SI, P30 females are significantly higher than P38 alone (Fig.3D). TH input is received from the VTA, which is adjacent to the substantia nigra compacta (SNc) which projects to other segments of the striatum. We quantified TH intensity in the VTA and SNc of males, finding a main effect of age where P30 shows increased TH density compared to P54 and no impact of SI (SFig.3.A-B). In females we see a main effect of age where P54 has significantly less TH than at P30, and a main effect of SI in the VTA and SNc where SI animals have overall increased TH (SFig.3C-D).

**Figure 3:**
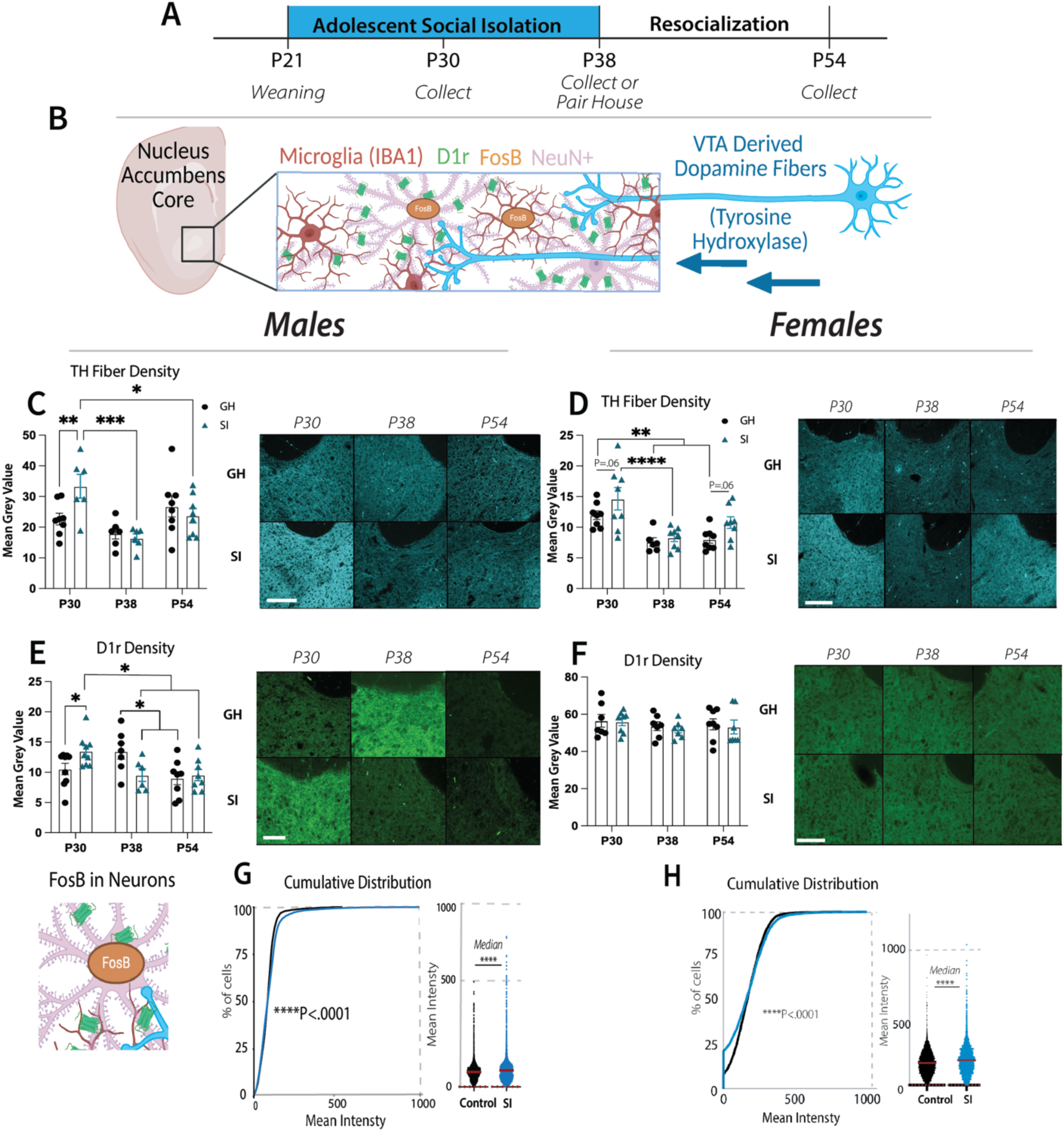
Adolescent SI Shifts TH Density and Activity in the NAc of Males and Females, D1r is Impacted Only in Males. A Experimental timeline showing collections during the SI period at P30, the end of the SI period at P38 or following rehousing at P54. B Schematic showing TH fiber input from the VTA to the NAc, D1rs on local SPNs and activity dependent FosB signaling in cell nuclei. C-F Dots represent single animal averages of mean intensity of either TH or D1r across 4 images per animal. Quantifications by Two-Way ANOVA with Tukey’s post hoc test followed by representative images where the scale bar equals 200µm. C NAc TH density quantification in males and D females. E NAc D1r density quantification in males and F females. G Cumulative istribution plot of FosB+ NeuN+ neurons at P30 followed by histogram, where each dot represents a cell in males and H females. ****P<.0001 Kolmogorov-Smirnov test for significant population distribution differences and Mann Whitney U test for differences in the median, N=6-8 animals 50-60 microglia 200-400 NeuN+ cells. All error bars = SEM, animals per age and condition are at least 4 distinct litters.

We next characterized NAc D1r development. In males we observe a significant interaction of age and SI where P30 SI males have significantly more D1r, while P38 SI males have significantly decreased D1r that normalizes at P54 (Fig.3E). In females we see no difference in D1r density due to SI (Fig.3F). We assessed other DA rich regions to ascertain whether the NAc is uniquely sensitive to the impacts of adolescent SI. We quantified D1r density in the dorsomedial striatum (DMS), finding no impact of SI in males or females (SFig.3E,G). Further in the NAc shell (NAcSh), a neighboring DA rich region that works in concert with the NAc^69^, there is no significant impact of SI in males or females (SFig.3F, H).

To assess activity changes in the NAc, we used IMARIS 3D reconstruction of NeuN and FosB+ intensity was quantified. In SI males at P30 there are more NeuN+ cells with higher FosB intensity, both in population distribution and population median (Fig.3G). In females there is a significant bimodal shift in NeuN+ FosB+ intensity population distribution and an SI-induced increase in the median (Fig.3H). Notably there are significant changes in FosB in NeuN+ and IBA+ microglia at P30, P38 and P54 in females (SFig.4F-J). These changes are not seen in males (SFig.4A-E).

Collectively in females we have found SI alters NAc TH input independent of D1r. SI also changes VTA and SNc TH, and activity dependent FosB NeuN+ and IBA+ microglia populations in females. In males we have found concurrent TH and D1r developmental shifts due to SI, with NAc activity differences observed in males at P30, the peak of microglia pruning. In males these data show a normalization of DA development of TH and D1r density at P54 following resocialization, while in females there is a trending increase in NAc TH at P54. There are sex specific impacts of SI with notable difference in DA development in males not present in females, consistent with sex specific mechanisms of microglia mediated DA system development. We next sought to characterize the impacts of adolescent SI on NAc microglia.

### 3.3 Adolescent SI persistently alters male microglial morphology, gene expression and increases engulfment of D1rs

Microglia morphology has been tied to alterations in function. Here we used a 2D sholl analysis approach to quantify microglia branch intersections as a readout for complexity across P30, P38 and P54 (Fig.4A-B). In the middle of the SI period at P30 there are no changes in microglia morphology in females (SFig.5G) or males (SFig.6B). In males at P38 there is a main effect of SI where microglia are hyper ramified showing increased branch intersections (Fig.4C). In P54 males there is a main effect of SI where microglia are hypo ramified showing decreased branch intersections (Fig.4D). Female SI microglia at P38 are hyper ramified, while at P54 they are hypo ramified (SFig.5A-B). These data show microglia across sexes are impacted by SI. We next sought to interrogate gene expression changes by microglia at P38 and P54 to infer functional alterations.

**Figure 4:**
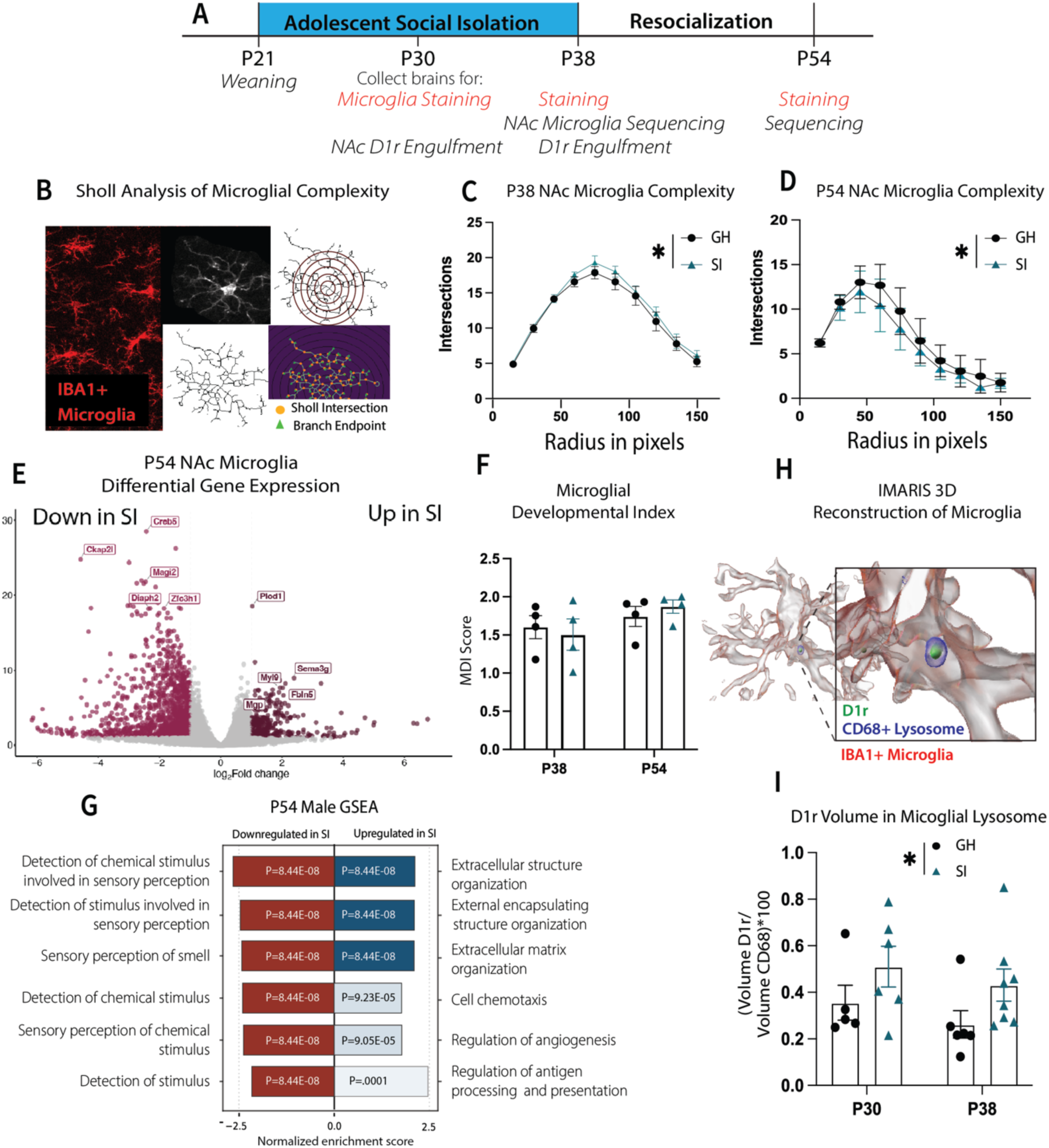
Adolescent SI Alters Male Microglia Morphology, Long Term Gene Expression and Increases Engulfment of NAc D1rs. A Experimental timeline showing experimental endpoints across SI study. B Schematic representation of sholl analysis to interrogate microglial complexity through use of concentric rings from the cell soma to quantify microglial branch complexity. C Microglial complexity at P38 and D P54, significance by two-way ANOVA where dots represent averages for the group (N=6-8 animals, 5-8 microglia per animal). E Differential gene expression using DESeq2 N=4 per group, genes shown as significant passed a threshold of padj<0.05 and LogFC>1. F Developmental index score on gene expression of P38 and P54 microglia, *ns by two-way ANOVA. G Gene Set Enrichment Analysis (GSEA) of P54 DEGs. H Graphical representation of IMARIS 3D reconstructed microglia and protein markers. I Quantification of engulfment of D1rs by microglia, where D1r volume is normalized to lysosome volume (CD68). Dots represent animal averages across 4 images, significance by two-way ANOVA *P<.05. All error bars = SEM, animals per age and condition are from at least 4 distinct litters.

To do this we macro dissected bilateral NAc/ventral striatum, isolating microglia from the tissue for bulk RNA-seq. In males at P38 there are 3 significantly different genes (SFig.6A), and at P54 more than 3400 including SI increases of complement genes *C1qa*, *C1qb* and *C1qc* (Fig.4E). In females there are 13 DEGs at P38 and 64 at P54 (SFig.5A-B). Early life stress has been shown to accelerate maturation^70^. We compared our gene set to a microglia maturation index to interrogate how SI might alter microglial transcriptional development^71^. We observed no differences as a result of SI or age between P38 and P54 male (Fig.4F) or female microglia (SFig.5E). These data reflect that microglia at P38 and P54 are not developmentally distinguishable via gene expression programs.

We next sought to characterize the functional changes in the observed gene expression changes via GSEA analysis. In SI males at P54 there is downregulation of sensory perception genes and upregulation of extracellular matrix, angiogenesis and antigen presentation coding genes (Fig.4G). In females we see an upregulation in transcriptional programs using RNA polymerase, neuron projection and axon guidance, postsynaptic chemical transmission (SFig.5F). In females there is downregulation of metabolic genes regulating mitochondrial function including NADH dehydrogenase complex assembly and ATP synthesis coupled electron transport (SFig.5F). Here we see sex-specific changes in microglia gene expression programs and ramification across development as a result of SI. There were no changes in maturation by gene expression, so we next determined if there are microglia number changes induced by SI within regions across development. We stained for IBA1, a homeostatic microglia signaling marker for a calcium binding protein that has been shown to be altered in developmental disorders including schizophrenia^72,73^.

Looking in females across development at P30, P38 and P54 we observe no SI-induced changes in microglia cell number in the NAc, DMS, NAcSh or VTA (SFig.5H,J,K,L,M). In females there is consistently an age-related decline in cell number from P30 to P38 into P54 across each of these brain regions (SFig.5H,J,K,L,M). There is a main effect of SI resulting in decreased overall NAc IBA1 intensity (SFig.5I), with trending effects in the DMS (p=0.1) and NAcSh (p=0.09) (SFig.5K,M). In males there is a significant interaction of SI and age on total NAc microglia number, where P30 GH animals have more microglia than at P54 (SFig.6C). In SI, there are significantly more microglia at P30 compared to P38 and P54, showing a steeper developmental decline (SFig.6C). There is a significant interaction effect on NAc IBA1 intensity showing a significant decline in SI males from P30 to P38 into P54, with no GH changes (SFig.6D). In the DMS there is an interaction effect on overall microglia number, where P54 SI animals have significantly fewer microglia than GH (SFig.6J), while there is no change in IBA1 intensity (SFig.6K). In the NAcSh of males there is an interaction effect of SI and age on total cell number (SFig.6L). At P30 SI animals have significantly more microglia than at P38 and P54, and more microglia than P30 GH animals (SFig.6L). In the VTA there is also an interaction effect where P30 SI animals have significantly more microglia than at P38 and P54, while in GH there are more microglia at P30 than P54 (SFig.6N). VTA IBA1 signaling shows a significant increase in P30 SI animals compared to P38 and P54, with no GH changes (SFig.6O). Collectively these data show sex and brain region specific developmental effects of SI. In males SI impacts microglia cell number across multiple brain regions, while females show little change in these endpoints. These data are consistent with immune hypotheses of developmental disorders where there is a male sex bias, as with autism and schizophrenia^74–77^.

Given the role of NAc microglia in regulating social behavior via developmental D1r engulfment, and the data reflecting NAc DA system and microglia changes, we used IMARIS 3D reconstruction to characterize NAc microglia features including engulfment/phagocytosis of D1rs (Fig.4A,H). We see no differences in overall lysosome marker CD68 intensity (SFig.6E p=0.1), area (SFig.6F), or normalized volume within microglia (SFig.6G). These data suggest there is not an overall change in phagocytic capacity within SI microglia. There is an interaction with individual microglia size, where SI microglia at P38 are larger than P30 with no differences in GH (SFig.6H). These data are consistent with SI-induced hyper ramification observed at P38 (Fig.4C). There is an age effect of overall microglia volume, showing a developmental increase from P30 to P38 (SFig.6I). Total volume increases, whereas total number does not (SFig.6C). We quantified phagocytosis of D1rs through measuring total volume of D1r in microglia lysosomes. We find a main effect of SI where there is an increase in D1r engulfment by NAc microglia in males (Fig.4H). This shows SI microglia play a more active role in shaping NAc DA signaling. This effect is not driven by overall number or volume of microglia. These data support the hypothesis that microglia may play a role in SI-induced behavioral changes. We next wanted to know what might be driving increased engulfment of D1rs by microglia.

### 3.4 DA activity alone does not regulate microglial engulfment of D1rs

There is evidence that microglia can sense and respond to DA activity^41^. In other brain regions, complement pruning by microglia has been shown to be neuronal activity dependent^63,78^. Here we sought to test the role of VTA derived DA signaling in NAc D1r density and microglia interactions. We tested this using a chemogenetic strategy in TH cre rats, where we selectively targeted VTA DA neurons with an excitatory (Gq), inhibitory (Gi) or control mCherry DREADD virus (Fig.5A). We injected the virus at P21, and began once daily i.p. 1mg/kg CNO injections for five days from P27-31. Animals were collected one hour following the final CNO injection. We observed robust reporter expression in the VTA and NAc at P31 following P21 injection (SFig.7A). The inhibitory DREADD was further validated using a cocaine-stimulated activity approach at P31, where CNO binding in Gi animals significantly decreased cocaine induced locomotion (SFig.7B-C). Quantification of NAc TH density showed no differences across virus conditions (Fig.5B). Quantification of NAc D1r density also showed no differences across virus condition (Fig.5C). Interrogation of FosB+ cells of the NAc showed no differences in total number (Fig.5D). Volume of individual microglia by IMARIS 3D reconstruction is unchanged (Fig.5E), and there is also no change in engulfed D1r using this chemogenetic strategy (Fig.5F). In summary, we find no evidence that VTA-DA activity into the NAc alone is sufficient to change microglial engulfment of D1rs. Next, we sought to uncover the role of microglial over-engulfment of D1rs after SI on behavior.

**Figure 5:**
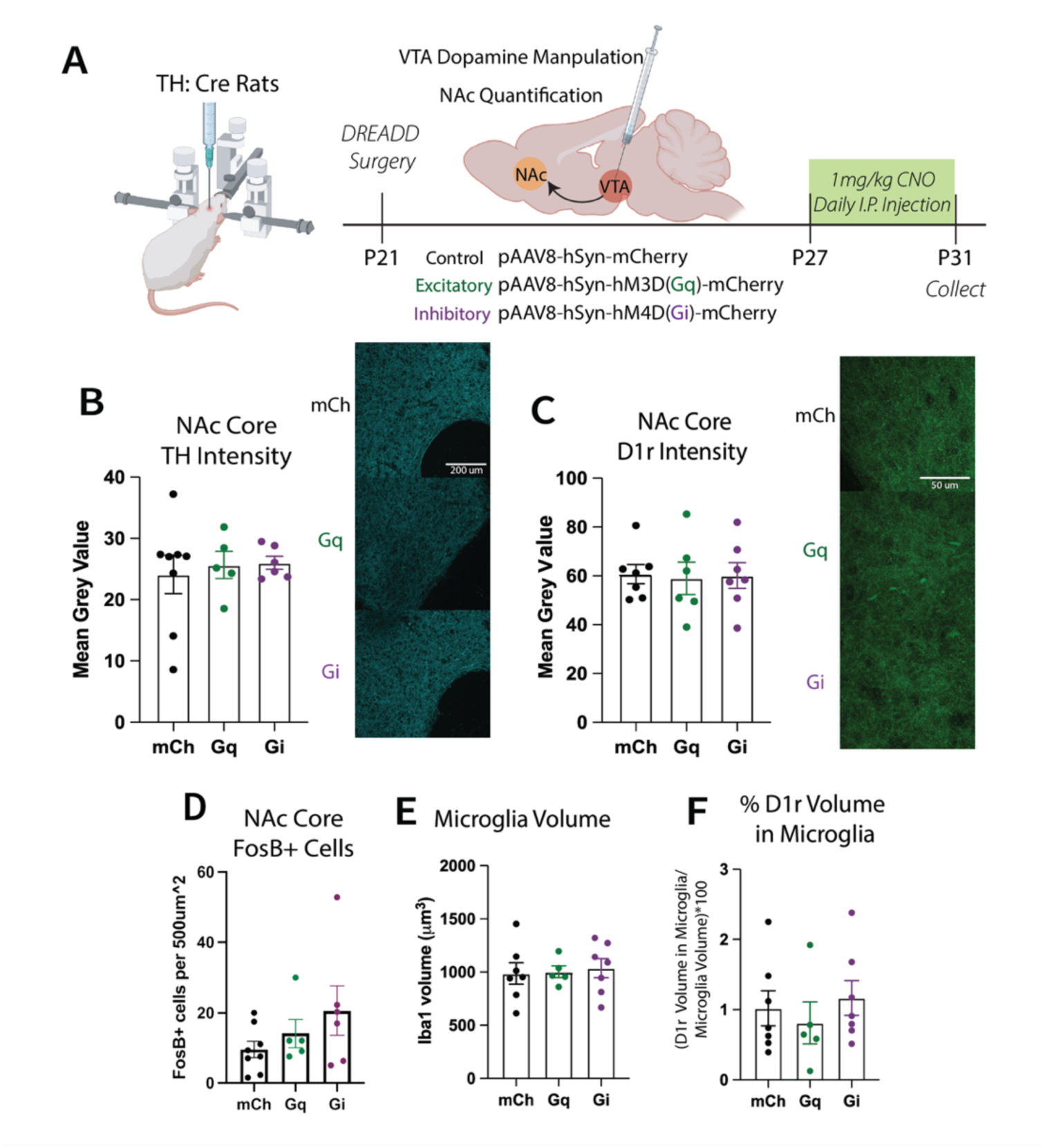
Chemogenetic Manipulation of the VTA During Adolescence was Insufficient to Alter NAc DA system Development in Males. A Experimental schematic showing the use of TH:cre rats and cre dependent DREADD expressing viruses to specifically target dopaminergic neurons of the VTA. Designer drug is administered once daily for 5 days, animals are collected one hour following final injection. Quantifications in the NAc to observe downstream impacts of circuit manipulation on microglia dopamine system interactions. B TH intensity quantification by one-way ANOVA followed by representative images across mCh control, Gq excitatory DREADD and Gi inhibitory DREADD where the scale bar is 200µm. Dots represent animal averages across 4 images, N=5-8. C D1r intensity quantification by one-way ANOVA followed by representative images where the scale bar is 50µm. D Quantification of FosB+ cells in the NAc normalized to total area by one-way ANOVA. E Quantification of volume of microglia by 3D reconstruction, N=5-7 with 5 microglia quantified per animal. E Volume of D1r inside of microglia, normalized to microglia volume quantified by one-way ANOVA, N=5-7 with 5 microglia quantified per animal. All error bars = SEM, animals are from at least 4 distinct litters.

### 3.5 NIF treatment interacts with social condition to co-regulate play and aggressive like behaviors in association with NAc D1r density

To test the role of microglia engulfment in SI behavioral expression, we used NIF—a hookworm protein that binds to the C3r subunit CD11b preventing complement engulfment^44^. We have previously demonstrated NIF injected to the NAc at P30 increases D1r density via preventing typical complement mediated engulfment^44^. Here, we bilaterally injected NIF (Fig.6A) or PBS control to NAc of P30 SI and GH males. All animals underwent play behavior testing at P38 and were collected within 2 hours. We observed a main effect of NIF and SI on NAc D1r density as expected (Fig.6B-C). In GH PBS animals, NAc D1r density is positively correlated to play behaviors, which is lost in GH NIF animals (Fig.6D, SFig.8A). In SI animals, in both NIF and PBS-treated groups, there is no correlation between NAc D1r density and play behaviors (Fig.6D, SFig.8A). We know SI increases aggressive like behaviors, including biting. To understand the relationship between NAc D1r and aggression we normalized bite time to total play time, correlating this to NAc D1r density across groups. NAc D1r density has a significant positive correlation to bite time in GH animals and a significant negative relationship in SI (SFig.8B). NIF-treated animals across social conditions show a positive correlation between NAc D1r density and normalized bite time (SFig.8C). These data implicate microglial function across social conditions in the regulation of bite behavior. Looking at group interactions, GH NIF NAc D1r density shows a significant positive correlation to bouncy gait, suggesting a hyperexcitability. GH NIF D1r density aloo shows a significant positive correlation to normalized bite time, while SI PBS animals a negative correlation (Fig.6E). NIF significantly shifted the relationship of NAc D1r density and bite behavior in GH and SI animals (Fig.6E). These data show that blocking complement-mediated engulfment by microglia with NIF interacts with social condition to regulate play and biting behavior. Moreover, these data suggest that microglial pruning is necessary for the functional coupling of NAc D1rs and behavioral output.

**Figure 6:**
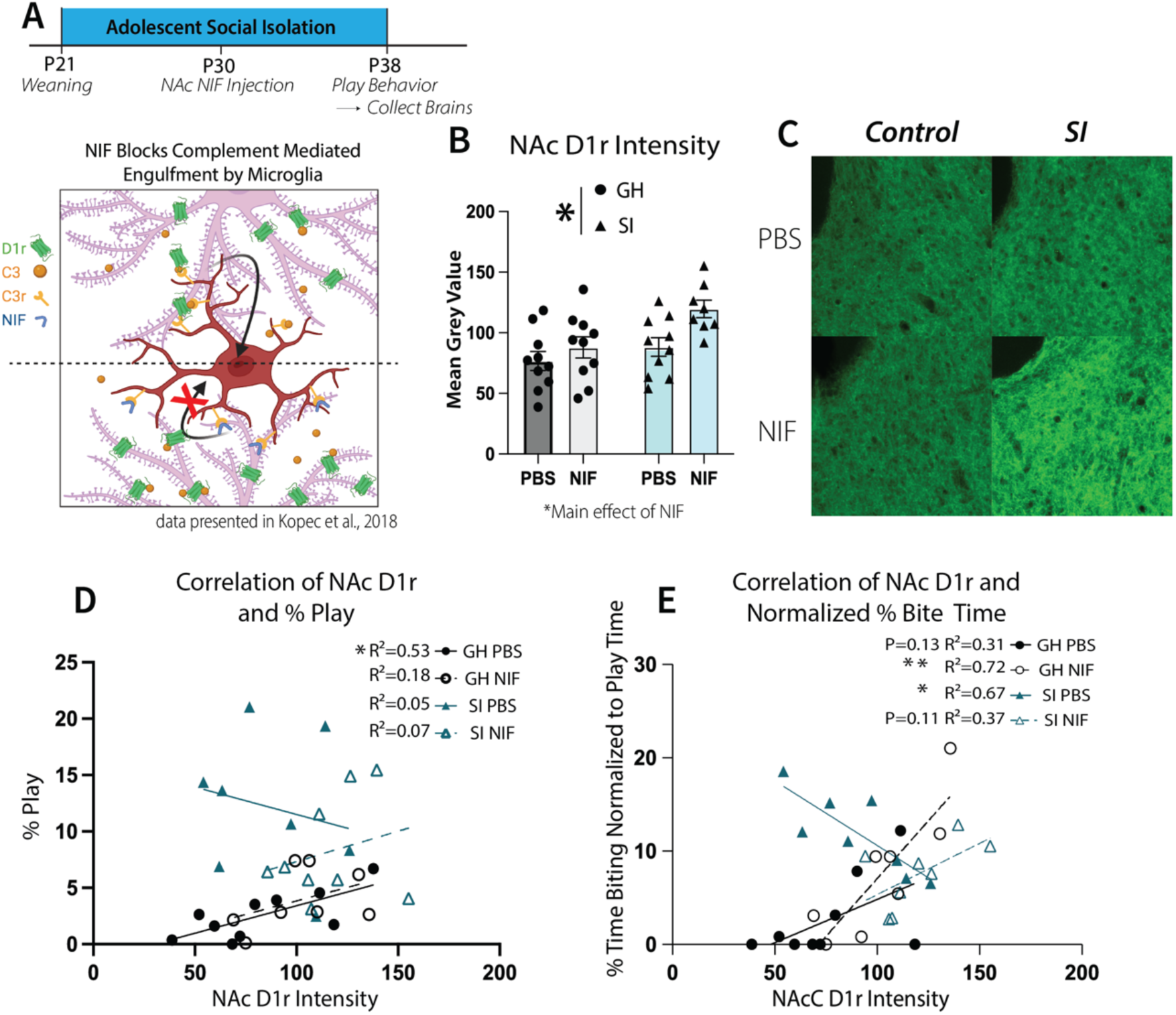
Social Experience and Complement Mediated Pruning by Microglia Interact to Direct Association of NAc D1r Density and Behavior. A Experimental schematic outlining the injection of NIF or PBS control to the NAc at P30, and later play behavior at P38. NIF blocks complement mediated engulfment by blocking the function of C3r. Brains are collected the same day following the 15 minute play bout. B NAc D1r quantification by two-way ANOVA *P<.05, dots represent animal averages of 4 images each, N=8-11 across 4 litters. Error bars = SEM. C Representative images of NAc D1r intensity across social condition and injectable. D PCC of NAc D1r density and % time spent playing within animals, regression lines for each group with R squared values represented. E PCC of NAc D1r density and normalized bite time with regression lines for each group plotted. N=8-10 *P<.05.

Next we investigated overall differences in play behavior repertoire due to treatment and social condition (Fig.7A,B). We performed PCA analysis of behaviors and represented this in 3D space capturing 70.85% of behavioral variability, where SI animals once again show more behavioral complexity space than GH animals (Fig.7C). Quantification of PC1, comprising primarily play behaviors, showed a main effect of SI (SFig.9A). PC2 made up of non-play social and affiliative behaviors revealed an interaction effect where SI NIF animals had greater scores than SI PBS and GH NIF (p=0.054) animals (SFig.9B). This was recapitulated in the data through quantification of total % time engaging in non-play social behaviors showing an interaction effect with a significant increase in SI NIF animals over SI PBS and GH NIF animals (Fig.7D). Quantification of play behavior shows SI NIF animals have significantly lower total % play compared to SI PBS, with no differences between SI NIF and GH animals (Fig.7E). This SI NIF decrease in play is primarily driven by interaction effects, impacting boxing, pinning, chasing and biting behaviors (SFig.10B,C,D,G,L,M). Given the interaction effect of SI and NIF on NAc D1r correlation to bite behavior (Fig.6E) and overall behavior engagement (SFig.10L,M), we sought to further understand the impacts of NIF on aggressive like behaviors in these animals.

**Figure 7:**
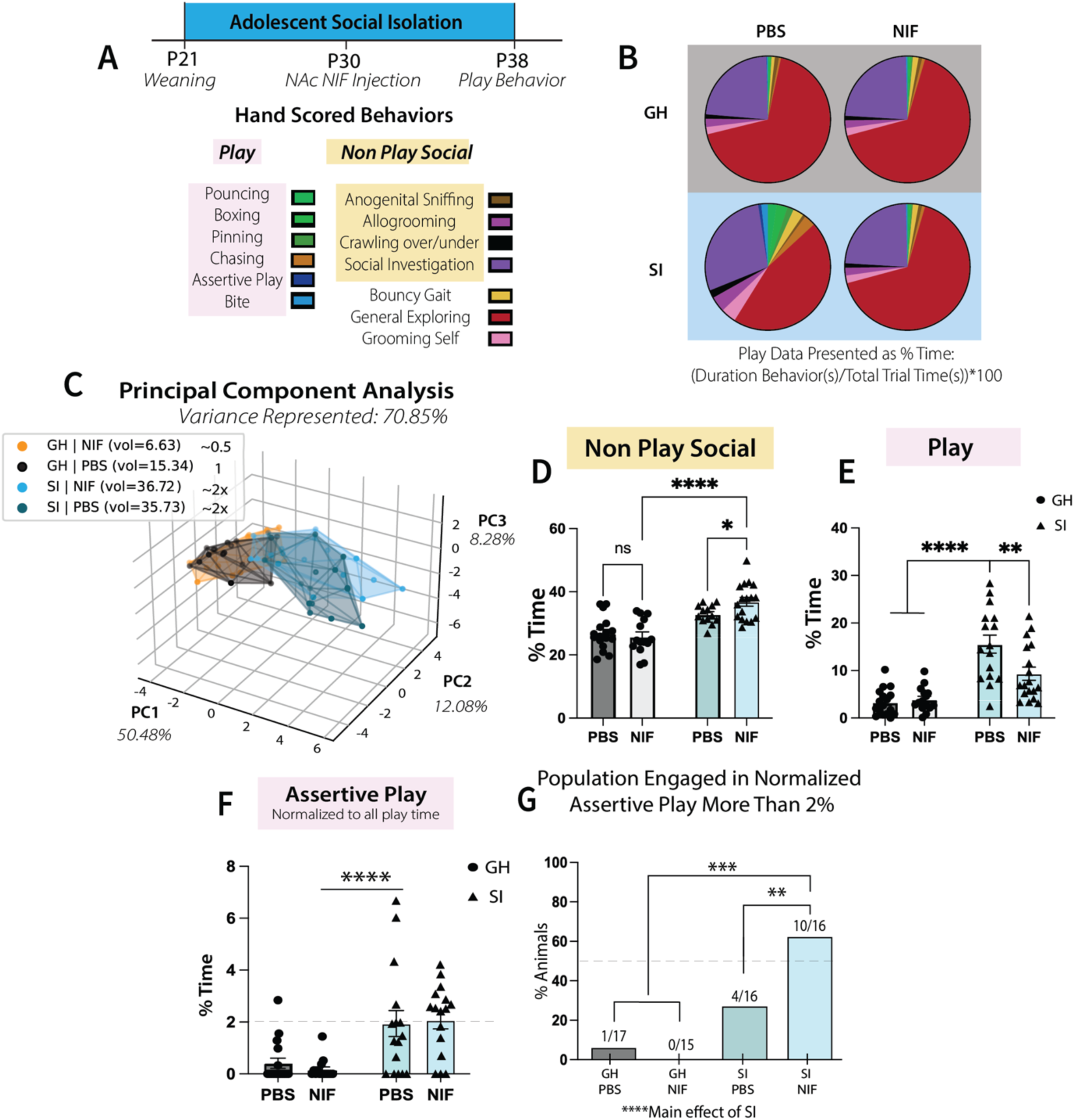
Social Experience and Microglia Complement Mediated Pruning Interact Regulating Affiliative and Aggressive like Behaviors. A Experimental timeline of NAc NIF or PBS injection at P30 with play behavior at P38, and associated hand scored behaviors. B Pie chart representation of behavioral repertoire. C PCA analysis of behavioral complexity represented in volumetric space with 70.85% of the variance captured. D Quantification of total non-play social behaviors, E play behaviors and F Assertive play normalized to all play time. G Quantification of animals engaged in normalized assertive play more than 2%, D-G analyses by two-way ANOVA with Tukey’s pos hoc test **P<.01,***<.001,****<.0001. N=15-18. Error bars = SEM. Total animals in each group are represented from at least 4 separate litters.

Percent bite time shows an interaction effect, where SI NIF animals have a trending decrease (p=0.064) compared to SI PBS, while total bite number shows a significant decrease in SI NIF compared to SI PBS animals (S10L,M). When looking at normalized bite time, and the percent of animals that engage it longest relative to total play time (1%> total play time) there is a main effect of NIF (SFig.10O). Interestingly there is no interaction effect of NIF and SI on assertive play, suggesting increased aggressive behavior that does not scale with bite (SFig.10K). Quantification of assertive play to total play time showed a main effect of SI (Fig.7F). Investigating the percent of animals engaged in normalized assertive play the most (2%> total play time) shows an interaction effect of SI and NIF, where significantly more SI NIF animals engage in high levels of assertive play compared to any other group (Fig.7G). Notably NIF does not increase assertive play in GH animals (Fig.7G). These data show microglia play a role in preventing exaggerated aggressive behavior in SI.

## 4. Discussion

Here we describe sex-specific impacts of an adolescent specific period of SI on brain development and behavior. In SI males and females we observe increased behavioral complexity including increased play and aggressive-like behaviors. There are persistent social changes in males after SI, specifically increased social novelty preference following a period of resocialization. In males both NAc TH and D1r developmental trajectory are shifted as a result of SI, while TH alone is altered by SI in females. We hypothesized microglia may mediate the observed male D1r differences, regulating SI-induced behavioral dysfunction. Indeed, in the NAc of SI males, microglia over engulf D1rs in adolescence. Microglial complement-driven pruning mediates the positive correlation of NAc D1rs to play behaviors in GH animals. Microglial pruning also mediates the negative correlation of NAc D1rs to biting. NIF interacts with SI, decreasing overall play and exaggerating non-reciprocal, aggressive-like assertive play behavior. Together, we describe a novel role for microglia as critical mediators of social development across social conditions. The data yield insights into developmental disorders including schizophrenia and autism which share features of social deficits and immune etiologies^79,80^. Further, the work presented here has great relevance towards understanding the long-term developmental impacts of the COVID 19 pandemic.

### 4.1 Adolescent SI Increases Behavior Complexity at P38 and Male Social Novelty at P54

Many studies outline the impacts of indefinite isolation into adulthood which make it difficult to parse developmental specificity^52,81,82^. In our model animals are isolated during the adolescent period from weaning at P21 until P38, at which point SI animals are housed together. At P54 in SI males we observed increased social novelty preference. These data fit other studies SI that have observed increases in novelty and reward seeking behaviors^47,59^. Some studies of adolescent SI find increases in social preference^20^, while others do not^83^, highlighting the importance of thorough characterization within studies due to strain differences and difficulty reconciling alternate SI paradigms with findings.

Changes in social interaction may contribute to the early mortality observed in SI^19,84,85^, as SI may disrupt necessary features of typical interactions that drive long term affiliation^86,87^. For these reasons we focused on understanding changes in behavioral features resulting from SI. Anogenital sniffing behavior in rats has been linked to dominance, in that more dominant rats are less likely to engage in anogenital sniffing and engage in direct facial investigation^88^. Here we see normalized reductions in time spent anogenitally sniffing by SI animals across sexes, suggesting increased dominance. When looking at play and aggressive like behaviors there are increases in SI males and females recapitulating prior findings^17,48,89–91^. Unexpectedly we observed normalized bite time to total play was significantly increased in SI females and not males, showing bite scales with play time in males and not females; highlighting a unique sex difference in SI induced expression of aggression. Overall these data provide a robust contemporary map of behavioral repertoire use across the sexes and social conditions. Further work looking at the long-term health outcomes of adolescent SI, using behavior repertoire features as long-term health predictors could be very insightful.

### 4.2 SI Has Sex Specific Effects on DA System Development

Developmental stress can accelerate maturation of stress sensitive circuits that regulate fear and complex emotional processing, including the DA system^92,93^. SI is a powerful developmental stressor that has been shown to increase DA release to the NAc in models of adolescent specific SI^94–96^ as well as persistent postweaning SI^52,53^. Through staining for both TH and D1r we aimed to capture VTA axon terminals in the NAc and receiving D1rs on local spiny projection neurons in the NAc. In females, we observe SI alterations in NAc and VTA TH fiber density, with no change in NAc D1r density. This suggests there is a difference in the utility of these VTA derived fibers independent of DA signaling directly. There are subpopulations of TH+ VTA neurons that co-release glutamate^97^, which may account for observed NAc TH changes, independent of D1r.

Differences in TH fiber density suggest there may be alterations in activity of these neurons. We used FosB as a proxy of activity, as FosB is an activity dependent marker whose splice variants are implicated in the transcriptional maintenance of neuronal gene programs that contribute to addiction liability^66^. We found a female specific increase of FosB in NAc neurons at all ages we investigated (P30, P38, P54). There is evidence for sex specificity in adolescent SI-induced excitability where there is increased presynaptic glutamate transmission from the hippocampus to the NAc in female mice, which may underlie differences seen here^96^. Surprisingly we also found increased microglial FosB at P30, P38 and P54. While FosB has been demonstrated to be expressed by microglia, and shown to regulate excitotoxic complement protein C5ar1/2 expression^98^, little else is known about the role of FosB in microglia.

In males we observe SI induced increases in both D1r and TH density in the NAc at P30, while there is significantly decreased D1r density at P38 in SI. This is in part mediated by SI microglia who over engulf NAc D1rs at P30 and P38. SI induced microglial over engulfment of D1rs shows microglia are significantly impacted by SI and alter how social circuits are sculpted in development. These are exciting new data demonstrating the environmental sensitivity of microglia in circuit refinement, with implications on mechanisms of psychiatric disease and addiction liability^26^.

We also observed an increase of FosB at P30 in NeuN+ cells of the NAc showing a neuronal activity is impacted during a period of increased engulfment by microglia. The concurrent change in TH and D1r suggests a change in dopamine signaling. We know microglia can prune synapses in an activity dependent manner^63^, to further understand the mechanism that drives D1r engulfment in SI, we tested whether DA activity was sufficient to modulate D1r engulfment by male microglia.

### 4.3 In SI Males Microglia Mediate NAc DA Developmental Shift in a DA-Independent Manner

We show that chemogenetic manipulation of VTA DA neurons is insufficient to drive D1r pruning or DA system changes with a daily CNO dose across the period of microglia pruning at P27-31 in males. Activity dependent pruning by microglia was first characterized using forskolin to stimulate activity and tetrodotoxin to inhibit activity from the eye to the lateral geniculate nucleus^42^. Here we use a chemogenetic approach using CNO as the designer drug. There is evidence that CNO back metabolizes into clozapine, a DA system agonist potentially clouding its utility in manipulating the DA system^99^. Further, people have used doses of 5mg/kg to 20mg/kg in rats to manipulate the system, making our dose of 1mg/kg modest in comparison^99,100^. Additionally the VTA is unique in that it is tonically active^97^; while CNO stays in the system for hours, there is still much of each day that would be unimpacted by chemogenetic manipulation. Taken together, in our hands we show that a consistent daily shift in VTA DA signaling is insufficient to alter NAc TH fiber density, D1r density or microglia engulfment of D1rs. These data suggest that DA activity alone is not driving SI-induced increases in D1r pruning by microglia. SI has been shown to induce glutamatergic signaling changes and circuit wide disinhibition^96^. The NAc receives top down excitatory glutamatergic inputs from the mPFC ^36,101,102^. Glutamate signaling to the NAc may be worth investigating to drive microglia pruning differences observed in SI. Ultimately, we sought to characterize the role of NAc microglial D1r engulfment on SI behavioral expression.

### 4.4 Microglia Integrate Social Experience and Sculpt NAc D1rs Shaping Behavior

The DA system encodes for social behaviors including play and aggression, as well as reward and aversion^23,24,103^. We find exciting data that show microglia play an intersectional role regulating play and aggression in the context of SI. We find a positive correlation of NAc D1r to play behavior in GH PBS animals only. Blocking complement-mediated microglial engulfment with NIF prevented this positive correlation, as did SI, showing that both microglia and social experience are necessary in NAc D1r:play associations. Preventing microglial complement pruning with NIF significantly altered D1r association to aggressive behaviors. NIF prevented the positive D1r:bite correlation in GH PBS animals, as well as the typically negative correlation in SI PBS animals. In NIF animals regardless of social history, D1r has a positive correlation to % time biting. These data show microglia complement pruning is directed by adolescent social experience to tune D1rs both play and biting behaviors.

Interestingly we found a SI x NIF interaction effect where assertive play is exaggerated in SI NIF animals. We see this interaction as a result of a decrease in overall play exhibited by SI NIF animals, without an associated decrease in overall assertive play. When normalizing assertive play to total play time, we see a higher proportion of animals engaging in high levels of assertive play. We observe that SI NIF animals engage in greater amounts of assertive play showing that microglia prevent exaggerated aggressive responses in the context of SI. To our knowledge this is the first study to outline this interaction of NAc D1r density, microglia and affiliative and aggressive behavior expression.

Our data fit with data in male mice showing NAc D1r x stress interactions altering social avoidance^104^. These data translate to a human PET study in a healthy population which found limbic striatal D1r to be positively correlated with social desirability/conformity and negatively associated with aggression^105^. Microglia have been shown to regulate social behaviors including aggression via proinflammatory cytokine signaling in response to immune activation^106,107^. Here we show microglia directly regulate developmental D1r density in the NAc to direct play and aggressive behavior in an experience-dependent manner. Collectively these data provide insight into the molecular mechanisms of social homeostasis, which s seeks to resolve how social drives are balanced across molecular neurobiological levels and social context^108^. We provide novel data implicating microglia at the crossroads of integrating social experience and shaping social behavior expression while patterning the developing DA system.

We know that engulfment of D1rs by microglia regulate social play in a D1r-dependent manner^44^, in the study presented here we do not know if it is D1r specifically driving these aggressive effects or other non-engulfed components coregulated with D1r by microglia. There is significant evidence for D1r involvement in aggressive behaviors making D1r a prime candidate^48,109,110^. Still, microglia play a significant role in regulating play and aggressive like behaviors in the context of SI. There remain many exciting questions to be answered; what drives microglial pruning in SI? and how might immune activation in response to infection or disease interact with SI-DA system interactions in development? Are compelling future directions for this research.

## Conclusions

Loneliness was declared an epidemic by the world health organization in 2023, following COVID 19 social distancing measures where 26 million adolescents experienced a period of social isolation^9,111,112^. We do not well understand the long term developmental consequences of social isolation (SI) on the brain and behavior. We know SI disrupts typical social behavior, while both autism and schizophrenia share features of disrupted social behavior^79,113^. Autism and schizophrenia also a share male sex bias, social deficits, immune origins and exhibit excitation/inhibition imbalances^80,82,113,114^. Here we use a rodent model of SI where we observe increased play and aggressive like behaviors^48,91^. We demonstrate in males a clear role for NAc microglia in coregulating play and aggressive like behaviors via D1r remodeling in a social experience dependent manner. We show complement mediated engulfment by microglia prevents exaggerated aggression in SI. Blocking microglial engulfment across social conditions drives biting behaviors. Microglia health and function in the face of social stress may be a key target toward mitigating negative outcomes of adolescent SI.

## Supporting information

Supplemental Video 1 Reciprocal Play

Supplemental Video 2 Assertive Play

## Acknowledgements

This work is supported by the National Institutes of Health specifically the National Institute on Drug of Abuse 5R01DA055414-03 and 1F31DA057116. The researchers would like to thank the Duke DLAR animal care team.

## Bibliography

1. Narayan, A.J., Lieberman, A.F., and Masten, A.S. (2021). Intergenerational transmission and prevention of adverse childhood experiences (ACEs). Clin. Psychol. Rev. 85, 101997. 10.1016/j.cpr.2021.101997.

2. Chang, X., Jiang, X., Mkandarwire, T., and Shen, M. (2019). Associations between adverse childhood experiences and health outcomes in adults aged 18–59 years. PLOS ONE 14, e0211850. 10.1371/journal.pone.0211850.

3. Buimer, E.E.L., Brouwer, R.M., Mandl, R.C.W., Pas, P., Schnack, H.G., and Hulshoff Pol, H.E. (2022). Adverse childhood experiences and fronto-subcortical structures in the developing brain. Front. Psychiatry 13. 10.3389/fpsyt.2022.955871.

4. Felitti, V.J., Anda, R.F., Nordenberg, D., Williamson, D.F., Spitz, A.M., Edwards, V., Koss, M.P., and Marks, J.S. (1998). Relationship of childhood abuse and household dysfunction to many of the leading causes of death in adults. The Adverse Childhood Experiences (ACE) Study. Am. J. Prev. Med. 14, 245–258. 10.1016/s0749-3797(98)00017-8.

5. Ma, L., Sun, F., and Tang, Z. (2018). Social Frailty is Associated with Physical Functioning, Cognition, and Depression, and Predicts Mortality. J. Nutr. Health Aging 22, 989–995. 10.1007/s12603-018-1054-0.

6. Fulford, D., and Holt, D.J. (2023). Social Withdrawal, Loneliness, and Health in Schizophrenia: Psychological and Neural Mechanisms. Schizophr. Bull. 49, 1138–1149. 10.1093/schbul/sbad099.

7. Kafali, H.Y., Bildik, T., Bora, E., Yuncu, Z., and Erermis, H.S. (2019). Distinguishing prodromal stage of bipolar disorder and early onset schizophrenia spectrum disorders during adolescence. Psychiatry Res. 275, 315–325. 10.1016/j.psychres.2019.03.051.

8. Miller, G.E., Chen, E., and Parker, K.J. (2011). Psychological Stress in Childhood and Susceptibility to the Chronic Diseases of Aging: Moving Towards a Model of Behavioral and Biological Mechanisms. Psychol. Bull. 137, 959–997. 10.1037/a0024768.

9. Child Population by Age Group Statistics https://datacenter.aecf.org/data/tables/101-child-population-by-age-group.

10. Arnett, J. (1992). Reckless behavior in adolescence: A developmental perspective. Dev. Rev. 12, 339–373. 10.1016/0273-2297(92)90013-R.

11. Giron, A.P., Ciranka, S., Schulz, E., van den Bos, W., Ruggeri, A., Meder, B., and Wu, C.M. (2023). Developmental changes in exploration resemble stochastic optimization. Nat. Hum. Behav. 7, 1955–1967. 10.1038/s41562-023-01662-1.

12. Bonawitz, E., Denison, S., Gopnik, A., and Griffiths, T.L. (2014). Win-Stay, Lose-Sample: A simple sequential algorithm for approximating Bayesian inference. Cognit. Psychol. 74, 35–65. 10.1016/j.cogpsych.2014.06.003.

13. Pellis, S.M., and Pellis, V.C. (2017). What is play fighting and what is it good for? Learn. Behav. 45, 355–366. 10.3758/s13420-017-0264-3.

14. Smith, P.K., and StGeorge, J.M. (2023). Play fighting (rough-and-tumble play) in children: Developmental and evolutionary perspectives. Int. J. Play 12, 113–126. 10.1080/21594937.2022.2152185.

15. Pellis, S.M., Field, E.F., Smith, L.K., and Pellis, V.C. (1997). Multiple Differences in the Play Fighting of Male and Female Rats. Implications for the Causes and Functions of Play. Neurosci. Biobehav. Rev. 21, 105–120. 10.1016/0149-7634(95)00060-7.

16. Vanderschuren, L.J., Niesink, R.J., and Van Ree, J.M. (1997). The neurobiology of social play behavior in rats. Neurosci. Biobehav. Rev. 21, 309–326. 10.1016/s0149-7634(96)00020-6.

17. Vanderschuren, L.J.M.J., Achterberg, E.J.M., and Trezza, V. (2016). The neurobiology of social play and its rewarding value in rats. Neurosci. Biobehav. Rev. 70, 86–105. 10.1016/j.neubiorev.2016.07.025.

18. Pellis, S.M., Pellis, V.C., Pelletier, A., and Leca, J.-B. (2019). Is play a behavior system, and, if so, what kind? Behav. Processes 160, 1–9. 10.1016/j.beproc.2018.12.011.

19. Yee, J.R., Cavigelli, S.A., Delgado, B., and McClintock, M.K. (2008). Reciprocal Affiliation Among Adolescent Rats During a Mild Group Stressor Predicts Mammary Tumors and Lifespan. Psychosom. Med. 70, 1050–1059. 10.1097/PSY.0b013e31818425fb.

20. Hol, T., Van den Berg, C.L., Van Ree, J.M., and Spruijt, B.M. (1999). Isolation during the play period in infancy decreases adult social interactions in rats. Behav. Brain Res. 100, 91–97. 10.1016/S0166-4328(98)00116-8.

21. Burgdorf, J., Kroes, R.A., and Moskal, J.R. (2017). Rough-and-tumble play induces resilience to stress in rats. NeuroReport 28, 1122. 10.1097/WNR.0000000000000864.

22. Oliveira, G. de F., Nguyen, A.T., Carreras-Simons, L., Niepsuj, T., Gadelhak, S.H., Johnson, A.K., Abdalla, A., Lev, E., Torres Román, S.G., Fuchs, S.N., et al. (2024). Puberty Blocker, Leuprolide, Reduces Sex Differences in Rough-and-Tumble Play and Anxiety-like Behavior in Juvenile Rats. Endocrinology 165, bqae046. 10.1210/endocr/bqae046.

23. Manduca, A., Servadio, M., Damsteegt, R., Campolongo, P., Vanderschuren, L.J., and Trezza, V. (2016). Dopaminergic Neurotransmission in the Nucleus Accumbens Modulates Social Play Behavior in Rats. Neuropsychopharmacol. Off. Publ. Am. Coll. Neuropsychopharmacol. 41, 2215–2223. 10.1038/npp.2016.22.

24. Volman, S.F., Lammel, S., Margolis, E.B., Kim, Y., Richard, J.M., Roitman, M.F., and Lobo, M.K. (2013). New Insights into the Specificity and Plasticity of Reward and Aversion Encoding in the Mesolimbic System. J. Neurosci. 33, 17569–17576. 10.1523/JNEUROSCI.3250-13.2013.

25. Crone, E.A., and Dahl, R.E. (2012). Understanding adolescence as a period of social–affective engagement and goal flexibility. Nat. Rev. Neurosci. 13, 636–650. 10.1038/nrn3313.

26. Chambers, R.A., Taylor, J.R., and Potenza, M.N. (2003). Developmental neurocircuitry of motivation in adolescence: A critical period of addiction vulnerability. Am. J. Psychiatry 160, 1041–1052. 10.1176/appi.ajp.160.6.1041.

27. Dow-Edwards, D., MacMaster, F.P., Peterson, B.S., Niesink, R., Andersen, S., and Braams, B.R. (2019). Experience during adolescence shapes brain development: From synapses and networks to normal and pathological behavior. Neurotoxicol. Teratol. 76, 106834. 10.1016/j.ntt.2019.106834.

28. Lewis, D.A. (1997). Development of the prefrontal cortex during adolescence: insights into vulnerable neural circuits in schizophrenia. Neuropsychopharmacol. Off. Publ. Am. Coll. Neuropsychopharmacol. 16, 385–398. 10.1016/S0893-133X(96)00277-1.

29. Bell, M.R. (2018). Comparing Postnatal Development of Gonadal Hormones and Associated Social Behaviors in Rats, Mice, and Humans. Endocrinology 159, 2596–2613. 10.1210/en.2018-00220.

30. Becht, A.I., Klapwijk, E.T., Wierenga, L.M., van der Cruijsen, R., Spaans, J., van der Aar, L., Peters, S., Branje, S., Meeus, W., and Crone, E.A. (2020). Longitudinal associations between structural prefrontal cortex and nucleus accumbens development and daily identity formation processes across adolescence. Dev. Cogn. Neurosci. 46, 100880. 10.1016/j.dcn.2020.100880.

31. Teicher, M.H., Krenzel, E., Thompson, A.P., and Andersen, S.L. (2003). Dopamine receptor pruning during the peripubertal period is not attenuated by NMDA receptor antagonism in rat. Neurosci. Lett. 339, 169–171. 10.1016/S0304-3940(02)01475-1.

32. Andersen, S.L., Thompson, A.T., Rutstein, M., Hostetter, J.C., and Teicher, M.H. (2000). Dopamine receptor pruning in prefrontal cortex during the periadolescent period in rats. Synapse 37, 167–169. 10.1002/1098-2396(200008)37:2%3C167::AID-SYN11%3E3.0.CO;2-B.

33. Andersen, S.L., Thompson, A.P., Krenzel, E., and Teicher, M.H. (2002). Pubertal changes in gonadal hormones do not underlie adolescent dopamine receptor overproduction. Psychoneuroendocrinology 27, 683–691. 10.1016/S0306-4530(01)00069-5.

34. Nardou, R., Lewis, E.M., Rothhaas, R., Xu, R., Yang, A., Boyden, E., and Dölen, G. (2019). Oxytocin-dependent reopening of a social reward learning critical period with MDMA. Nature 569, 116–120. 10.1038/s41586-019-1075-9.

35. Yoshida, J., Oñate, M., Khatami, L., Vera, J., Nadim, F., and Khodakhah, K. (2022). Cerebellar Contributions to the Basal Ganglia Influence Motor Coordination, Reward Processing, and Movement Vigor. J. Neurosci. 42, 8406–8415. 10.1523/JNEUROSCI.1535-22.2022.

36. Stolyarova, A., and Wikenheiser, A.M. (2020). Can the VTA Come Out to Play? Only When the mPFC’s Predictions Go Astray! Neuron 105, 593–595. 10.1016/j.neuron.2020.01.036.

37. Schultz, W., Dayan, P., and Montague, P.R. (1997). A neural substrate of prediction and reward. Science 275, 1593–1599. 10.1126/science.275.5306.1593.

38. Heydari, S., and Holroyd, C.B. (2016). Reward positivity: Reward prediction error or salience prediction error? Psychophysiology 53, 1185–1192. 10.1111/psyp.12673.

39. Berridge, K.C. (2007). The debate over dopamine’s role in reward: the case for incentive salience. Psychopharmacology (Berl.) 191, 391–431. 10.1007/s00213-006-0578-x.

40. de Jong, J.W., Afjei, S.A., Pollak Dorocic, I., Peck, J.R., Liu, C., Kim, C.K., Tian, L., Deisseroth, K., and Lammel, S. (2019). A Neural Circuit Mechanism for Encoding Aversive Stimuli in the Mesolimbic Dopamine System. Neuron 101, 133–151.e7. 10.1016/j.neuron.2018.11.005.

41. Badimon, A., Strasburger, H.J., Ayata, P., Chen, X., Nair, A., Ikegami, A., Hwang, P., Chan, A.T., Graves, S.M., Uweru, J.O., et al. (2020). Negative feedback control of neuronal activity by microglia. Nature 586, 417–423. 10.1038/s41586-020-2777-8.

42. Schafer, D.P., Lehrman, E.K., Kautzman, A.G., Koyama, R., Mardinly, A.R., Yamasaki, R., Ransohoff, R.M., Greenberg, M.E., Barres, B.A., and Stevens, B. (2012). Microglia sculpt postnatal neural circuits in an activity and complement-dependent manner. Neuron 74, 691–705. 10.1016/j.neuron.2012.03.026.

43. Webster, R., Taberner, J., Edgington, A., Guhan, S., Varghese, J., Feeney, H., Blocker, L., and Jezequel, S.G. (1999). Role of sialylation in determining the pharmacokinetics of neutrophil inhibitory factor (NIF) in the Fischer 344 rat. Xenobiotica Fate Foreign Compd. Biol. Syst. 29, 1141–1155. 10.1080/004982599238010.

44. Kopec, A.M., Smith, C.J., Ayre, N.R., Sweat, S.C., and Bilbo, S.D. (2018). Microglial dopamine receptor elimination defines sex-specific nucleus accumbens development and social behavior in adolescent rats. Nat. Commun. 9, 3769. 10.1038/s41467-018-06118-z.

45. Nie, X., Kitaoka, S., Tanaka, K., Segi-Nishida, E., Imoto, Y., Ogawa, A., Nakano, F., Tomohiro, A., Nakayama, K., Taniguchi, M., et al. (2018). The Innate Immune Receptors TLR2/4 Mediate Repeated Social Defeat Stress-Induced Social Avoidance through Prefrontal Microglial Activation. Neuron 99, 464–479.e7. 10.1016/j.neuron.2018.06.035.

46. Wang, J., Chen, H.-S., Li, H.-H., Wang, H.-J., Zou, R.-S., Lu, X.-J., Wang, J., Nie, B.-B., Wu, J.-F., Li, S., et al. (2023). Microglia-dependent excessive synaptic pruning leads to cortical underconnectivity and behavioral abnormality following chronic social defeat stress in mice. Brain. Behav. Immun. 109, 23–36. 10.1016/j.bbi.2022.12.019.

47. Arakawa, H. (2005). Interaction between isolation rearing and social development on exploratory behavior in male rats. Behav. Processes 70, 223–234. 10.1016/j.beproc.2005.07.002.

48. Takahashi, A. (2025). The role of social isolation stress in escalated aggression in rodent models. Neurosci. Res. 211, 75–84. 10.1016/j.neures.2022.07.009.

49. Tulogdi, Á., Tóth, M., Barsvári, B., Biró, L., Mikics, É., and Haller, J. (2014). Effects of resocialization on post-weaning social isolation-induced abnormal aggression and social deficits in rats. Dev. Psychobiol. 56, 49–57. 10.1002/dev.21090.

50. Bendersky, C.J., Milian, A.A., Andrus, M.D., De La Torre, U., and Walker, D.M. (2021). Long-Term Impacts of Post-weaning Social Isolation on Nucleus Accumbens Function. Front. Psychiatry 12.

51. Begni, V., Sanson, A., Pfeiffer, N., Brandwein, C., Inta, D., Talbot, S.R., Riva, M.A., Gass, P., and Mallien, A.S. (2020). Social isolation in rats: Effects on animal welfare and molecular markers for neuroplasticity. PLOS ONE 15, e0240439. 10.1371/journal.pone.0240439.

52. Karkhanis, A.N., Locke, J.L., McCool, B.A., Weiner, J.L., and Jones, S.R. (2014). Social isolation rearing increases nucleus accumbens dopamine and norepinephrine responses to acute ethanol in adulthood. Alcohol. Clin. Exp. Res. 38, 2770–2779. 10.1111/acer.12555.

53. Karkhanis, A.N., Leach, A.C., Yorgason, J.T., Uneri, A., Barth, S., Niere, F., Alexander, N.J., Weiner, J.L., McCool, B.A., Raab-Graham, K.F., et al. (2019). Chronic Social Isolation Stress during Peri-Adolescence Alters Presynaptic Dopamine Terminal Dynamics via Augmentation in Accumbal Dopamine Availability. ACS Chem. Neurosci. 10, 2033–2044. 10.1021/acschemneuro.8b00360.

54. Han, X., Wang, W., Xue, X., Shao, F., and Li, N. (2011). Brief social isolation in early adolescence affects reversal learning and forebrain BDNF expression in adult rats. Brain Res. Bull. 86, 173–178. 10.1016/j.brainresbull.2011.07.008.

55. Baarendse, P.J.J., Counotte, D.S., O’Donnell, P., and Vanderschuren, L.J.M.J. (2013). Early Social Experience Is Critical for the Development of Cognitive Control and Dopamine Modulation of Prefrontal Cortex Function. Neuropsychopharmacology 38, 1485–1494. 10.1038/npp.2013.47.

56. Hinton, E.A., Li, D.C., Allen, A.G., and Gourley, S.L. (2019). Social Isolation in Adolescence Disrupts Cortical Development and Goal-Dependent Decision-Making in Adulthood, Despite Social Reintegration. eNeuro 6, ENEURO.0318-19.2019. 10.1523/ENEURO.0318-19.2019.

57. Whitaker, L.R., Degoulet, M., and Morikawa, H. (2013). Social deprivation enhances VTA synaptic plasticity and drug-induced contextual learning. Neuron 77, 335–345. 10.1016/j.neuron.2012.11.022.

58. Fosnocht, A.Q., Lucerne, K.E., Ellis, A.S., Olimpo, N.A., and Briand, L.A. (2019). Adolescent Social Isolation Increases Cocaine Seeking in Male and Female Mice. Behav. Brain Res. 359, 589–596. 10.1016/j.bbr.2018.10.007.

59. Noschang, C., Lampert, C., Krolow, R., and de Almeida, R.M.M. (2021). Social isolation at adolescence: a systematic review on behaviour related to cocaine, amphetamine and nicotine use in rats and mice. Psychopharmacology (Berl.) 238, 927–947. 10.1007/s00213-021-05777-z.

60. Bordt, E.A., Block, C.L., Petrozziello, T., Sadri-Vakili, G., Smith, C.J., Edlow, A.G., and Bilbo, S.D. (2020). Isolation of Microglia from Mouse or Human Tissue. STAR Protoc. 1, 100035. 10.1016/j.xpro.2020.100035.

61. Gunaydin, L.A., Grosenick, L., Finkelstein, J.C., Kauvar, I.V., Fenno, L.E., Adhikari, A., Lammel, S., Mirzabekov, J.J., Airan, R.D., Zalocusky, K.A., et al. (2014). Natural Neural Projection Dynamics Underlying Social Behavior. Cell 157, 1535–1551. 10.1016/j.cell.2014.05.017.

62. Stephan, A.H., Barres, B.A., and Stevens, B. (2012). The complement system: an unexpected role in synaptic pruning during development and disease. Annu. Rev. Neurosci. 35, 369–389. 10.1146/annurev-neuro-061010-113810.

63. Schafer, D.P., Lehrman, E.K., Kautzman, A.G., Koyama, R., Mardinly, A.R., Yamasaki, R., Ransohoff, R.M., Greenberg, M.E., Barres, B.A., and Stevens, B. (2012). Microglia Sculpt Postnatal Neural Circuits in an Activity and Complement-Dependent Manner. Neuron 74, 691–705. 10.1016/j.neuron.2012.03.026.

64. Caillé, I., Dumartin, B., and Bloch, B. (1996). Ultrastructural localization of D1 dopamine receptor immunoreactivity in rat striatonigral neurons and its relation with dopaminergic innervation. Brain Res. 730, 17–31. 10.1016/0006-8993(96)00424-6.

65. Daubner, S.C., Le, T., and Wang, S. (2011). Tyrosine hydroxylase and regulation of dopamine synthesis. Arch. Biochem. Biophys. 508, 1–12. 10.1016/j.abb.2010.12.017.

66. Robison, A.J., and Nestler, E.J. (2022). ΔFOSB: A Potentially Druggable Master Orchestrator of Activity-Dependent Gene Expression. ACS Chem. Neurosci. 13, 296–307. 10.1021/acschemneuro.1c00723.

67. Cates, H.M., Lardner, C.K., Bagot, R.C., Neve, R.L., and Nestler, E.J. (2019). Fosb Induction in Nucleus Accumbens by Cocaine Is Regulated by E2F3a. eNeuro 6. 10.1523/ENEURO.0325-18.2019.

68. Nestler, E.J., Barrot, M., and Self, D.W. (2001). ΔFosB: A sustained molecular switch for addiction. Proc. Natl. Acad. Sci. 98, 11042–11046. 10.1073/pnas.191352698.

69. Bassareo, V., Musio, P., and Di Chiara, G. (2011). Reciprocal responsiveness of nucleus accumbens shell and core dopamine to food- and drug-conditioned stimuli. Psychopharmacology (Berl.) 214, 687–697. 10.1007/s00213-010-2072-8.

70. Bath, K.G., Manzano-Nieves, G., and Goodwill, H. (2016). Early life stress accelerates behavioral and neural maturation of the hippocampus in male mice. Horm. Behav. 82, 64–71. 10.1016/j.yhbeh.2016.04.010.

71. Hanamsagar, R., Alter, M.D., Block, C.S., Sullivan, H., Bolton, J.L., and Bilbo, S.D. (2018). Generation of a microglial developmental index in mice and in humans reveals a sex difference in maturation and immune reactivity. Glia 66, 460. 10.1002/glia.23277.

72. Hill, S.L., Shao, L., and Beasley, C.L. (2021). Diminished levels of the chemokine fractalkine in post-mortem prefrontal cortex in schizophrenia but not bipolar disorder. World J. Biol. Psychiatry Off. J. World Fed. Soc. Biol. Psychiatry 22, 94–103. 10.1080/15622975.2020.1755451.

73. Snijders, G.J.L.J., van Zuiden, W., Sneeboer, M.A.M., Berdenis van Berlekom, A., van der Geest, A.T., Schnieder, T., MacIntyre, D.J., Hol, E.M., Kahn, R.S., and de Witte, L.D. (2021). A loss of mature microglial markers without immune activation in schizophrenia. Glia 69, 1251–1267. 10.1002/glia.23962.

74. Bordt, E.A., Moya, H.A., Jo, Y.C., Ravichandran, C.T., Bankowski, I.M., Ceasrine, A.M., McDougle, C.J., Carlezon, W.A., and Bilbo, S.D. (2024). Gonadal hormones impart male-biased behavioral vulnerabilities to immune activation via microglial mitochondrial function. Brain. Behav. Immun. 115, 680–695. 10.1016/j.bbi.2023.11.010.

75. Robinson-Agramonte, M. de los A., Noris García, E., Fraga Guerra, J., Vega Hurtado, Y., Antonucci, N., Semprún-Hernández, N., Schultz, S., and Siniscalco, D. (2022). Immune Dysregulation in Autism Spectrum Disorder: What Do We Know about It? Int. J. Mol. Sci. 23, 3033. 10.3390/ijms23063033.

76. Napolitano, A., Schiavi, S., La Rosa, P., Rossi-Espagnet, M.C., Petrillo, S., Bottino, F., Tagliente, E., Longo, D., Lupi, E., Casula, L., et al. (2022). Sex Differences in Autism Spectrum Disorder: Diagnostic, Neurobiological, and Behavioral Features. Front. Psychiatry 13, 889636. 10.3389/fpsyt.2022.889636.

77. Pruessner, M., Cullen, A.E., Aas, M., and Walker, E.F. (2017). The neural diathesis-stress model of schizophrenia revisited: An update on recent findings considering illness stage and neurobiological and methodological complexities. Neurosci. Biobehav. Rev. 73, 191–218. 10.1016/j.neubiorev.2016.12.013.

78. Presumey, J., Bialas, A.R., and Carroll, M.C. (2017). Complement System in Neural Synapse Elimination in Development and Disease. Adv. Immunol. 135, 53–79. 10.1016/bs.ai.2017.06.004.

79. Hakenova, K., Mikulecka, A., Holubova, K., Chvojkova, M., Slamberova, R., Jurcovicova, J., Cechova, B., Ponist, S., Horacek, J., and Vales, K. (2025). A new two-hit animal model for schizophrenia research: Consequences on social behavior. IBRO Neurosci. Rep. 19, 38–49. 10.1016/j.ibneur.2025.05.012.

80. Doll, C.A., and Broadie, K. (2014). Impaired activity-dependent neural circuit assembly and refinement in autism spectrum disorder genetic models. Front. Cell. Neurosci. 8. 10.3389/fncel.2014.00030.

81. Lapiz, M.D.S., Fulford, A., Muchimapura, S., Mason, R., Parker, T., and Marsden, C.A. (2003). Influence of postweaning social isolation in the rat on brain development, conditioned behavior, and neurotransmission. Neurosci. Behav. Physiol. 33, 13–29. 10.1023/a:1021171129766.

82. Ayilara, G.O., and Owoyele, B.V. (2023). Neuroinflammation and microglial expression in brains of social-isolation rearing model of schizophrenia. IBRO Neurosci. Rep. 15, 31–41. 10.1016/j.ibneur.2023.05.010.

83. Kim, M., Netser, S., Wagner, S., and Harony-Nicolas, H. (2025). Juvenile social isolation in Sprague Dawley rats does not have a lasting impact on social behavior in adulthood. Sci. Rep. 15, 12981. 10.1038/s41598-025-95920-z.

84. Holt-Lunstad, J., Smith, T.B., and Layton, J.B. (2010). Social relationships and mortality risk: a meta-analytic review. PLoS Med. 7, e1000316. 10.1371/journal.pmed.1000316.

85. Hilakivi-Clarke, L., and de Oliveira Andrade, F. (2023). Social Isolation and Breast Cancer. Endocrinology 164, bqad126. 10.1210/endocr/bqad126.

86. Zhang, F., You, Z., Fan, C., Gao, C., Cohen, R., Hsueh, Y., and Zhou, Z. (2014). Friendship quality, social preference, proximity prestige, and self-perceived social competence: interactive influences on children’s loneliness. J. Sch. Psychol. 52, 511–526. 10.1016/j.jsp.2014.06.001.

87. Hyland, P., Shevlin, M., Cloitre, M., Karatzias, T., Vallières, F., McGinty, G., Fox, R., and Power, J.M. (2019). Quality not quantity: loneliness subtypes, psychological trauma, and mental health in the US adult population. Soc. Psychiatry Psychiatr. Epidemiol. 54, 1089–1099. 10.1007/s00127-018-1597-8.

88. Wesson, D.W. (2013). Sniffing Behavior Communicates Social Hierarchy. Curr. Biol. 23, 575–580. 10.1016/j.cub.2013.02.012.

89. Lee, C.R., Chen, A., and Tye, K.M. (2021). The neural circuitry of social homeostasis: Consequences of acute versus chronic social isolation. Cell 184, 1500–1516. 10.1016/j.cell.2021.02.028.

90. Einon, D.F., Humphreys, A.P., Chivers, S.M., Field, S., and Naylor, V. (1981). Isolation has permanent effects upon the behavior of the rat, but not the mouse, gerbil, or guinea pig. Dev. Psychobiol. 14, 343–355. 10.1002/dev.420140407.

91. Orsucci, I.C., Becker, K.D., Ham, J.R., Lee, J.D.A., Bowden, S.M., and Veenema, A.H. (2025). To play or not to play? Effects of social isolation length and stimulus rat familiarity on social play engagement in three laboratory rat strains. Physiol. Behav. 302, 115080. 10.1016/j.physbeh.2025.115080.

92. Gee, D.G., Gabard-Durnam, L.J., Flannery, J., Goff, B., Humphreys, K.L., Telzer, E.H., Hare, T.A., Bookheimer, S.Y., and Tottenham, N. (2013). Early developmental emergence of human amygdala–prefrontal connectivity after maternal deprivation. Proc. Natl. Acad. Sci. U. S. A. 110, 15638–15643. 10.1073/pnas.1307893110.

93. Manzano Nieves, G., Bravo, M., Baskoylu, S., and Bath, K.G. (2020). Early life adversity decreases pre-adolescent fear expression by accelerating amygdala PV cell development. eLife 9, e55263. 10.7554/eLife.55263.

94. McWain, M.A., Pace, R.L., Nalan, P.A., and Lester, D.B. (2022). Age-dependent effects of social isolation on mesolimbic dopamine release. Exp. Brain Res. 240, 2803–2815. 10.1007/s00221-022-06449-w.

95. Social isolation postweaning alters reward-related dopamine dynamics in a region-specific manner in adolescent male rats - ClinicalKey https://www-clinicalkey-com.proxy.lib.duke.edu/#!/content/playContent/1-s2.0-S235228952400016X?returnurl=https:%2F%2Flinkinghub.elsevier.com%2Fretrieve%2Fpii%2FS235228952400016X%3Fshowall%3Dtrue&referrer=https:%2F%2Fpubmed.ncbi.nlm.nih.gov%2F.

96. Deutschmann, A.U., Kirkland, J.M., and Briand, L.A. (2022). Adolescent social isolation induced alterations in nucleus accumbens glutamate signalling. Addict. Biol. 27, e13077. 10.1111/adb.13077.

97. Morales, M., and Margolis, E.B. (2017). Ventral tegmental area: cellular heterogeneity, connectivity and behaviour. Nat. Rev. Neurosci. 18, 73–85. 10.1038/nrn.2016.165.

98. Nomaru, H., Sakumi, K., Katogi, A., Ohnishi, Y.N., Kajitani, K., Tsuchimoto, D., Nestler, E.J., and Nakabeppu, Y. (2014). Fosb gene products contribute to excitotoxic microglial activation by regulating the expression of complement C5a receptors in microglia. Glia 62, 1284–1298. 10.1002/glia.22680.

99. Manvich, D.F., Webster, K.A., Foster, S.L., Farrell, M.S., Ritchie, J.C., Porter, J.H., and Weinshenker, D. (2018). The DREADD agonist clozapine N-oxide (CNO) is reverse-metabolized to clozapine and produces clozapine-like interoceptive stimulus effects in rats and mice. Sci. Rep. 8, 3840. 10.1038/s41598-018-22116-z.

100. MacLaren, D.A.A., Browne, R.W., Shaw, J.K., Krishnan Radhakrishnan, S., Khare, P., España, R.A., and Clark, S.D. (2016). Clozapine N-Oxide Administration Produces Behavioral Effects in Long–Evans Rats: Implications for Designing DREADD Experiments. eNeuro 3, ENEURO.0219-16.2016. 10.1523/ENEURO.0219-16.2016.

101. Choi, T.-Y., Jeon, H., Jeong, S., Kim, E.J., Kim, J., Jeong, Y.H., Kang, B., Choi, M., and Koo, J.W. (2024). Distinct prefrontal projection activity and transcriptional state conversely orchestrate social competition and hierarchy. Neuron 112, 611–627.e8. 10.1016/j.neuron.2023.11.012.

102. Pintori, N., Piva, A., Mottarlini, F., Díaz, F.C., Maggi, C., Caffino, L., Fumagalli, F., and Chiamulera, C. (2024). Brief exposure to enriched environment rapidly shapes the glutamate synapses in the rat brain: A metaplastic fingerprint. Eur. J. Neurosci. 59, 982–995. 10.1111/ejn.16279.

103. Dai, B., Zheng, B., Dai, X., Cui, X., Yin, L., Cai, J., Zhuo, Y., Tritsch, N.X., Zweifel, L.S., Li, Y., et al. (2025). Experience-dependent dopamine modulation of male aggression. Nature 639, 430–437. 10.1038/s41586-024-08459-w.

104. Burek, D.J., and Carlezon, W.A. (2026). Stress-induced adaptations in nucleus accumbens dopamine D1 receptor-expressing cells correspond to social avoidance behavior in male mice. IBRO Neurosci. Rep. 21, 335–341. 10.1016/j.ibneur.2026.07.009.

105. Plavén-Sigray, P., Matheson, G.J., Gustavsson, P., Stenkrona, P., Halldin, C., Farde, L., and Cervenka, S. (2018). Is dopamine D1 receptor availability related to social behavior? A positron emission tomography replication study. PLoS ONE 13, e0193770. 10.1371/journal.pone.0193770.

106. Takahashi, A. (2024). Associations of the immune system in aggression traits and the role of microglia as mediators. Neuropharmacology 256, 110021. 10.1016/j.neuropharm.2024.110021.

107. Kopec, A.M., Smith, C.J., and Bilbo, S.D. (2019). Neuro-Immune Mechanisms Regulating Social Behavior: Dopamine as Mediator? Trends Neurosci. 42, 337–348. 10.1016/j.tins.2019.02.005.

108. Matthews, G.A., and Tye, K.M. (2019). Neural mechanisms of social homeostasis. Ann. N. Y. Acad. Sci. 1457, 5–25. 10.1111/nyas.14016.

109. Breese, G.R., Criswell, H.E., and Mueller, R.A. (1990). Evidence that lack of brain dopamine during development can increase the susceptibility for aggression and self-injurious behavior by influencing D1-dopamine receptor function. Prog. Neuropsychopharmacol. Biol. Psychiatry 14 Suppl, S65–80. 10.1016/0278-5846(90)90089-y.

110. Golden, S.A., Jin, M., Heins, C., Venniro, M., Michaelides, M., and Shaham, Y. (2019). Nucleus Accumbens Drd1-Expressing Neurons Control Aggression Self-Administration and Aggression Seeking in Mice. J. Neurosci. 39, 2482–2496. 10.1523/JNEUROSCI.2409-18.2019.

111. Office of the Surgeon General (OSG). (2023). Our Epidemic of Loneliness and Isolation. Wash. DC US Dep. Health Hum. Serv.

112. Sikali, K. (2020). The dangers of social distancing: How COVID-19 can reshape our social experience. J. Community Psychol. 48, 2435–2438. 10.1002/jcop.22430.

113. Fan, G., Ma, J., Ma, R., Suo, M., Chen, Y., Zhang, S., Zeng, Y., and Chen, Y. (2023). Microglia Modulate Neurodevelopment in Autism Spectrum Disorder and Schizophrenia. Int. J. Mol. Sci. 24, 17297. 10.3390/ijms242417297.

114. Imamura, A., Morimoto, Y., Ono, S., Kurotaki, N., Kanegae, S., Yamamoto, N., Kinoshita, H., Tsujita, T., Okazaki, Y., and Ozawa, H. (2020). Genetic and environmental factors of schizophrenia and autism spectrum disorder: insights from twin studies. J. Neural Transm. 127, 1501–1515. 10.1007/s00702-020-02188-w.

