## Supplemental Method and Figures for "Microglia sculpt dopamine receptor-mediated adolescent behaviors in a social experience dependent manner"

**Supplementary Method: Inhibitory DREADD Validation Experiment**

Animals underwent stereotaxic surgeries targeting the VTA as described in the primary text of this manuscript. To validate the inhibitory DREADD, Gi or mCherry control animals underwent open field testing at P30 following a 20mg/kg i.p. cocaine injection. Animals were allowed to habituate to the room for 30 minutes before undergoing a 30 minute baseline trial in the open field chamber. Animals were then given an i.p. 1mg/kg CNO injection and returned to the chamber for 30 minutes. Animals were then given a 20mg/kg i.p. injection of cocaine and returned to the open field where they were recorded for 75 more minutes. All animals were collected 15 minutes following the trial.

Supplementary Figures

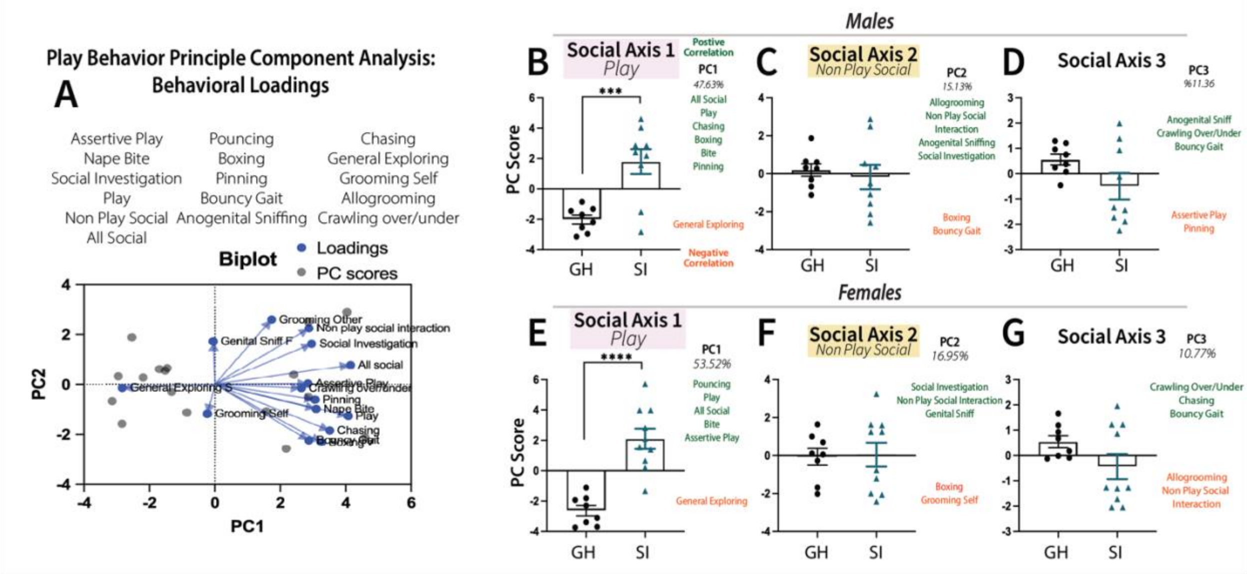


SF1: Adolescent SI Behavioral Repertoire Breakdown. A-I Quantification of behaviors, represented as total percent of trial across males and females. A Chasing, B pouncing, C Boxing, D Bite, D’ % Bite time normalized to play, E Assertive play, F crawling over/under, G anogenital sniffing, G’ % Anogenital sniff normalized to total non-play social time, H grooming and I bouncy gait.. *P<.05***<.001****<.0001 by multiple unpaired T-Test N=8-10. Error bars = SEM.

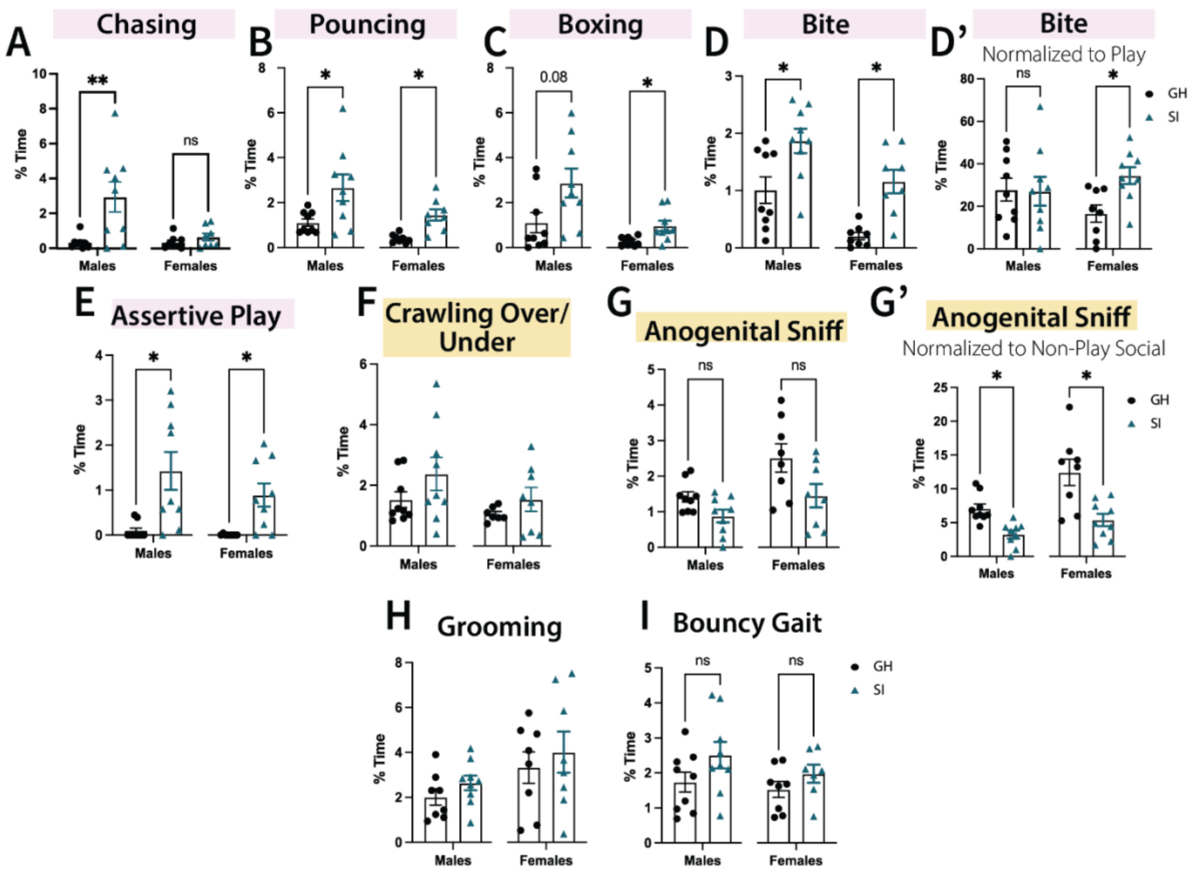


SF2: PCA Analysis of Play Behavior. A Behavioral loadings for principal component analysis and biplot representation of loadings in 2D behavioral space. B-D Comparisons of PC1, social axis 1-play, PC2, social axis 2-non play social and PC3 other social/affiliative behaviors in males and E-G females. Significance by multiple unpaired T-Test N=8-10. Error bars = SEM.***<.001****<.0001.

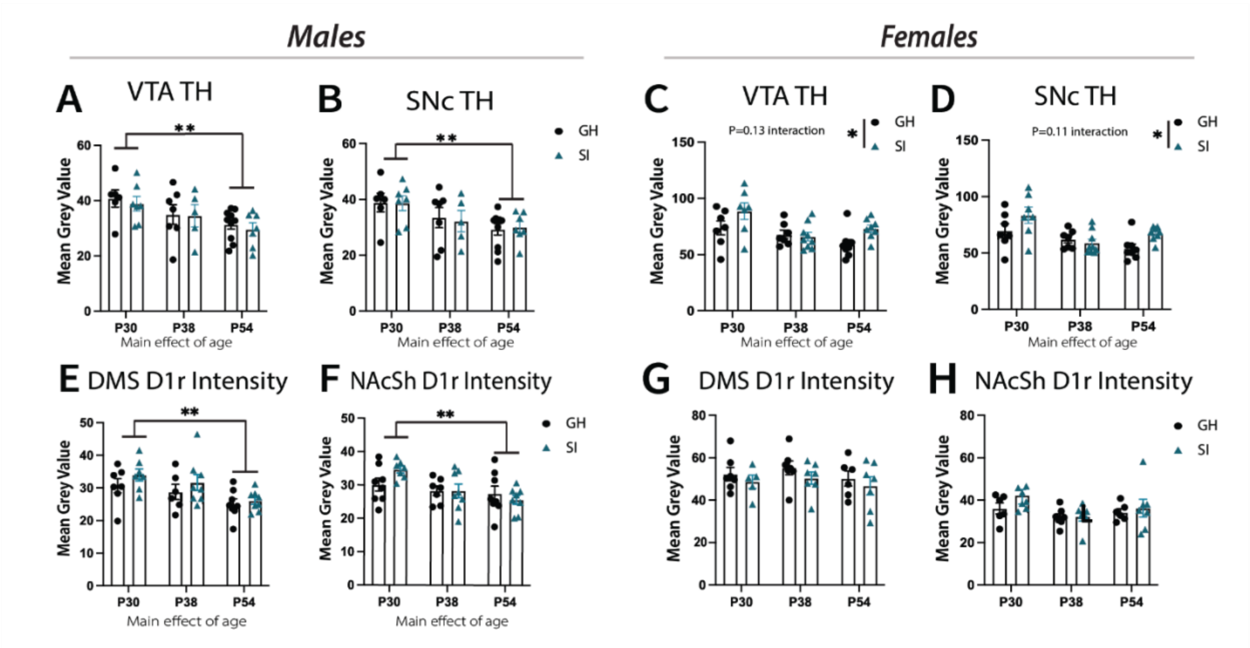


SF3: Adolescent SI Alters Mesolimbic Development in a Sex Specific Manner. Dots represent animal averages across 2-4 images quantified for all endpoints in this figure N= 6-9 across 4 litters for males and females. A Quantification of TH fiber density in the VTA and B SNc of males. B Quantification of VTA TH fiber density and C SNc TH fiber density in females. D1r density quantification in the E DMS and F NAcSh of males. G DMS and H NAcSh D1r density in females. Quantifications by two-way ANOVA with Tukey's post hoc testing, *P<.05,**<.01***<.001****<.0001.

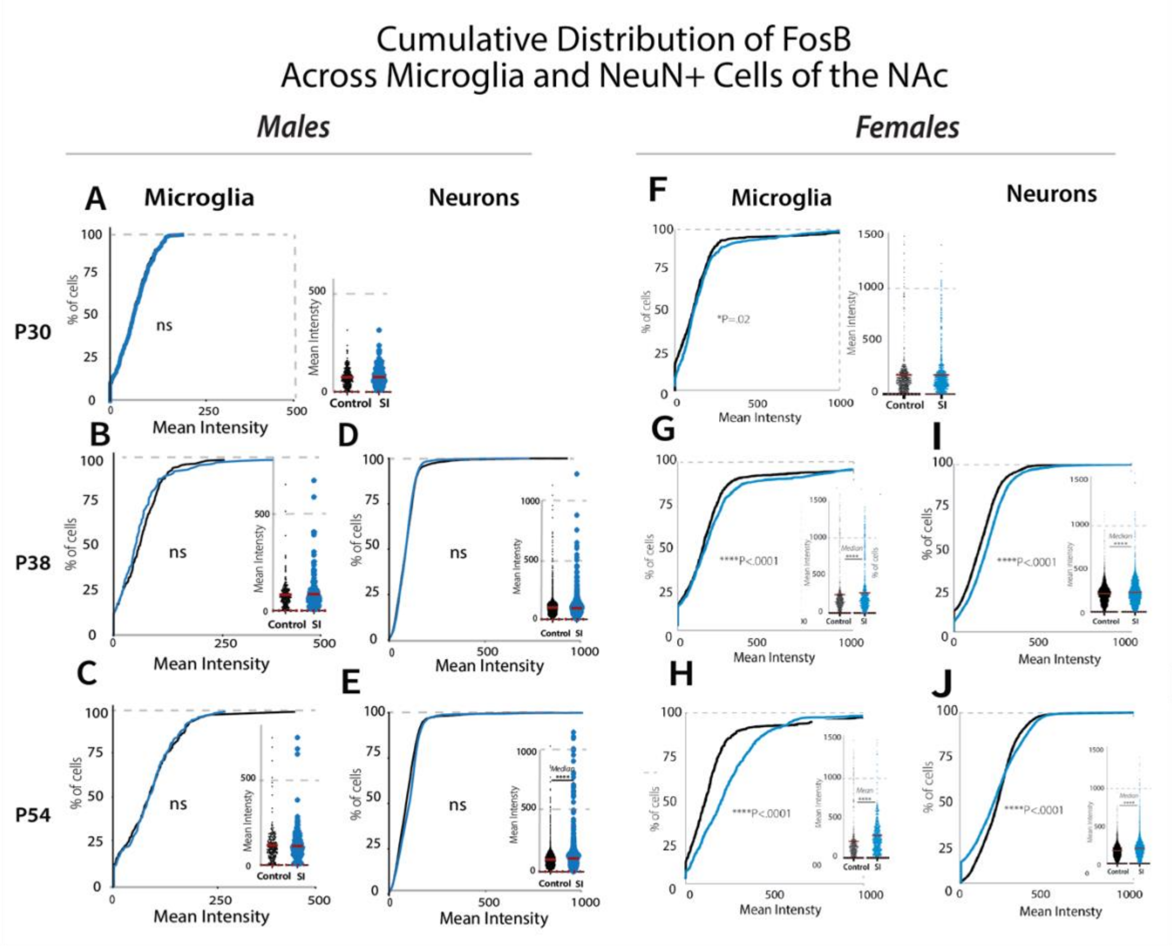


SF4: Developmental Characterization of Activity Dependent FosB in Microglia and Neurons. A Cumulative distribution plot of FosB+ IBA+ male NAc microglia at P30 followed by histograms where each dot represents a cell. B this same quantification in P38 and C P54 microglia. D Cumulative distribution and histogram of FosB+ NeuN+ cells at P38 and E P54. F-J quantifications are in females. F Cumulative distribution and histogram of FosB+ microglia at P30, G P38 and H P54. . I Quantification of FosB+ neurons at P38 and J P54. Significance by Kolmogorov-Smirnov test for significant population differences in distribution and Mann-Whitney U test for population differences in the median, ****$P<.0001$ N=6-8 animals 50-60 microglia 200-400 NeuN+ cells across 2-3 images. All error bars = SEM, animals per age and condition are at least 4 distinct litters.

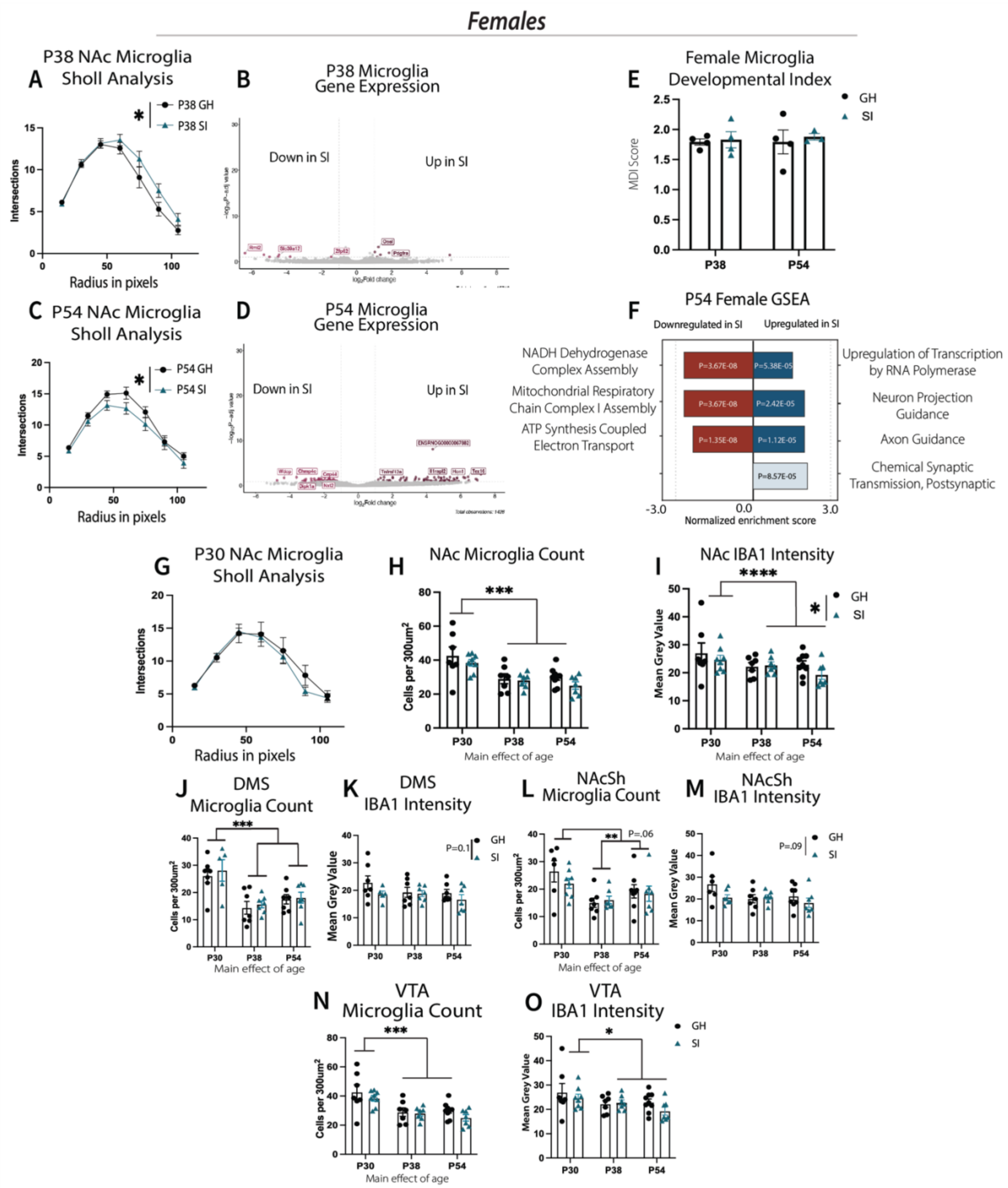


SF5: Female Specific Differences in NAc Microglia Features Including Morphology, Number and Gene Expression. A P38 microglia sholl analysis by two-way ANOVA. Dots represent group average, 5-8 microglia per animal (N=7-8). B P38 microglia differential gene expression using DESeq2 N=4 per group, genes shown as significant passed a threshold of padj<0.05 and LogFC>1. C P54 microglia sholl analysis and D gene expression, N=3-4. E Quantification of

microglia developmental index at P38 and P54. F Gene set enrichment analysis (GSEA) of P54 female DEGs. G P30 microglia sholl analysis, ns. For H-O dots represent animal averages of quantifications across 2-4 images per animal (N=5-9). H NAc microglia cell count and I IBA1 intensity. J DMS microglia cell count and K IBA1 intensity. L NAcSh Microglia cell count and M IBA1+ intensity. N VTA Microglia count and O IBA1 intensity. Quantifications by two-way ANOVA with Tukey's post hoc testing, *P<.05,**<.01***<.001****<.0001, all error bars represent SEM.

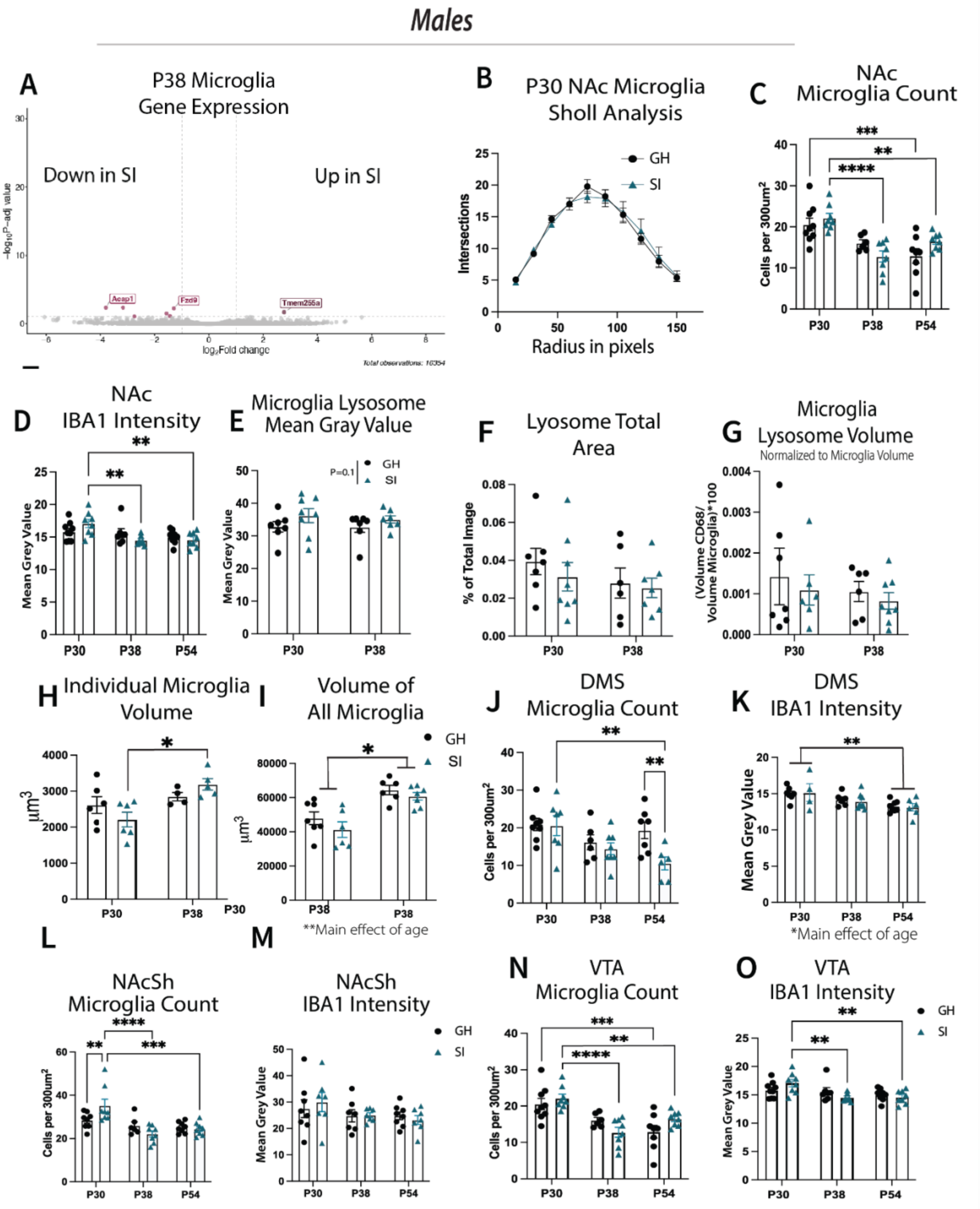


SF6: Male Specific Differences in Microglia Features in the NAc, and SI Altered Microglia Development in Other DA Rich Regions. B Sholl analysis of NAc microglial complexity at P30, ns by two-way ANOVA. Dots represent group average, 5-8 microglia per animal. C Microglia cell body count per 300μm$^2$, dots represent animal averages (N=7-9) measured across 4 images. D

IBA1 intensity quantified across 4 images. E Microglia lysosome, CD68, intensity. F Total thresholded area of CD68 signal. G IMARIS reconstructed lysosome volume normalized to microglia volume. H IMARIS quantification of individual microglia volume, dots represent animal averages of 5 microglia per animal. I volume all microglia in the total volume space of 2-3 Z-stack images per animal, N=5-9. J DMS microglia cell count, K DMS IBA1 intensity. L NAcSh microglia cell count, M NAcSh IBA1 intensity. N VTA Microglia cell count, N VTA IBA1 intensity. C-O, significance by two-way ANOVA with Tukey's post hoc unless otherwise specified. *P<.05,**<.01, ***<.001, ****<.0001, all error bars represent SEM.

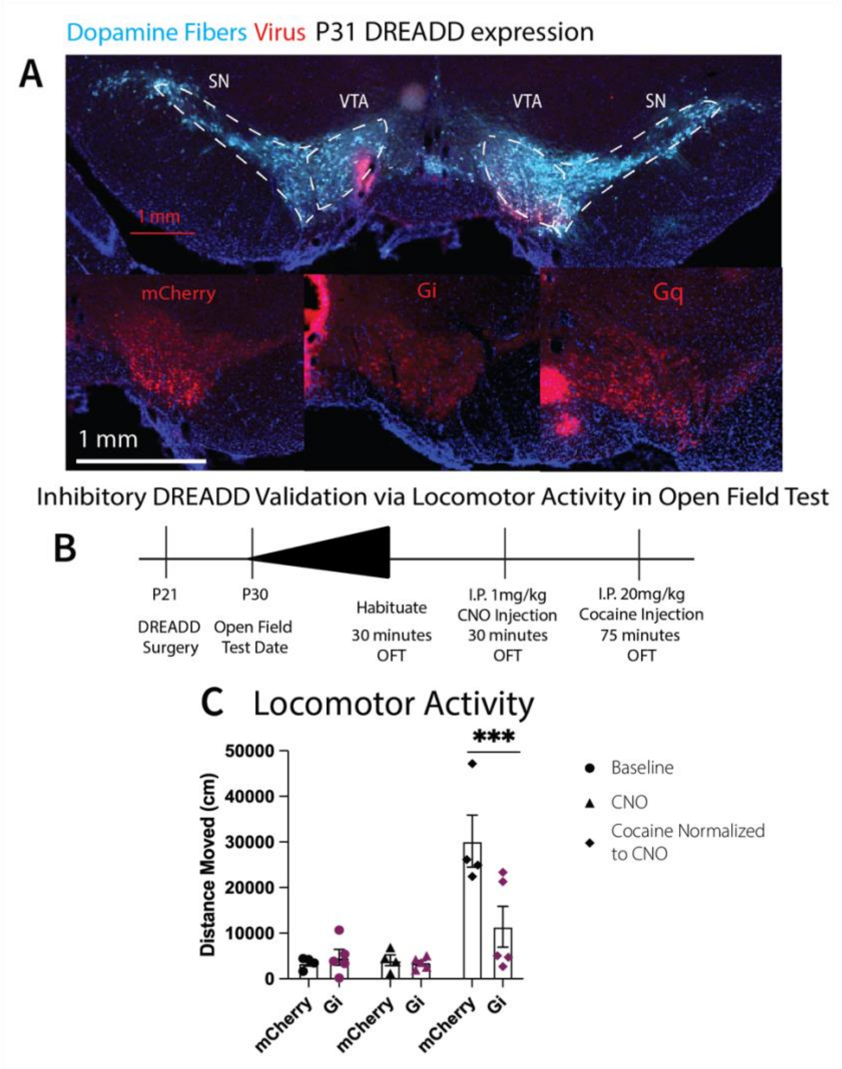


SF7: DREADD Expression at P31 and Inhibitory Behavioral Validation. A Representative image of overlapping TH and viral expression in the VTA and expression of the different DREADDs at P31. B Experimental setup where animals receive viral injection of an inhibitory DREADD to the NAc at P21, and undergo open field testing (OFT) at P30. On the day of testing animals undergo a 30 minute baseline test, followed by 1 mg/kg CNO injection and another 30 minutes OFT. Finally animals receive 20 mg/kg cocaine and undergo 75 minutes of OFT. C Quantification of locomotor activity N=4-5, significance by two-way ANOVA and Tukey's post hoc test, ***P<.001.

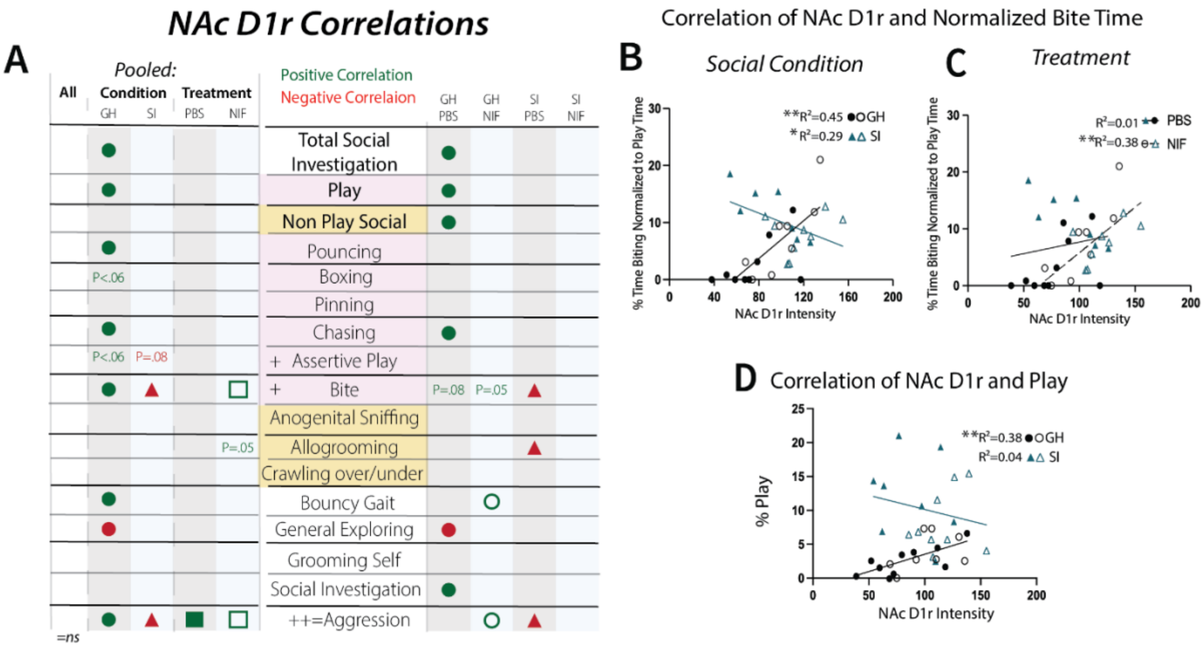


SF8: NIF and SI Interact Altering D1r:Behavior Correlates. A Correlations of NAc D1r to behavior, collapsed across all groups or pooled by social condition and injectable. GH animals are circles, SI animals triangles. PBS treated animals are closed shapes while NIF treated animals are open. Symbol presence denotes significance by PCC *P=<.05. B Correlation of NAc D1r and bite time by social condition and C injectable. D Correlation of NAc D1r and play across social condition, N=17-18 per social condition and injectable.

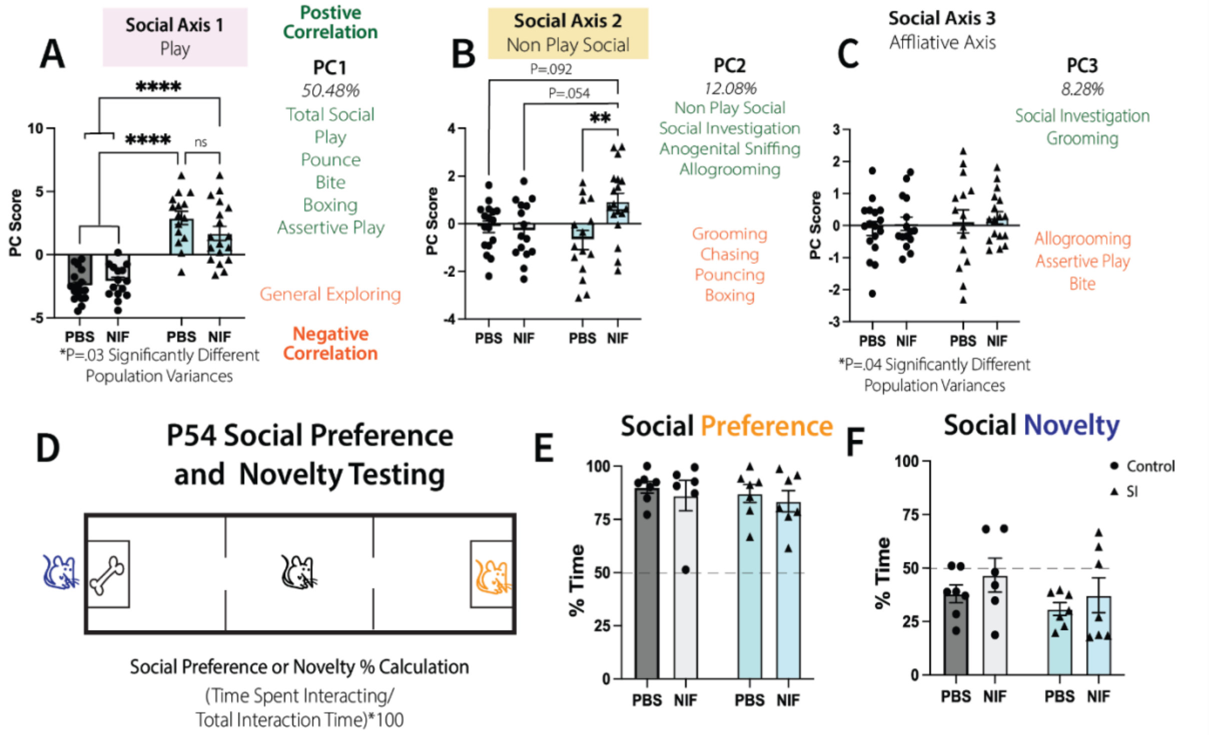


SF9: PCA Quantification of SI and NIF Interacting on Behavior. A PCA analysis PC1/social axis 1 quantification, capturing play behaviors. B PC2/social axis 3 quantification capturing largely non play social behaviors. C PC3/social axis 3 quantification capturing affiliative behaviors, dots represent single animals N=16-19. Significance by two-way ANOVA with Tukey's post hoc test. D PCC of normalized allogrooming and biting behavior, regression line plotted with $R^2$ values represented. E PCC of biting and assertive play behavior, N=8-10. F Social preference and novelty testing setup at P54, where there are five minutes spent on the preference portion and 5 minutes on the novelty portion. G Social preference and H social novelty scores, N=6-7. *P<.05, **<.01, ***<.001, ****<.0001. Error bars = SEM.

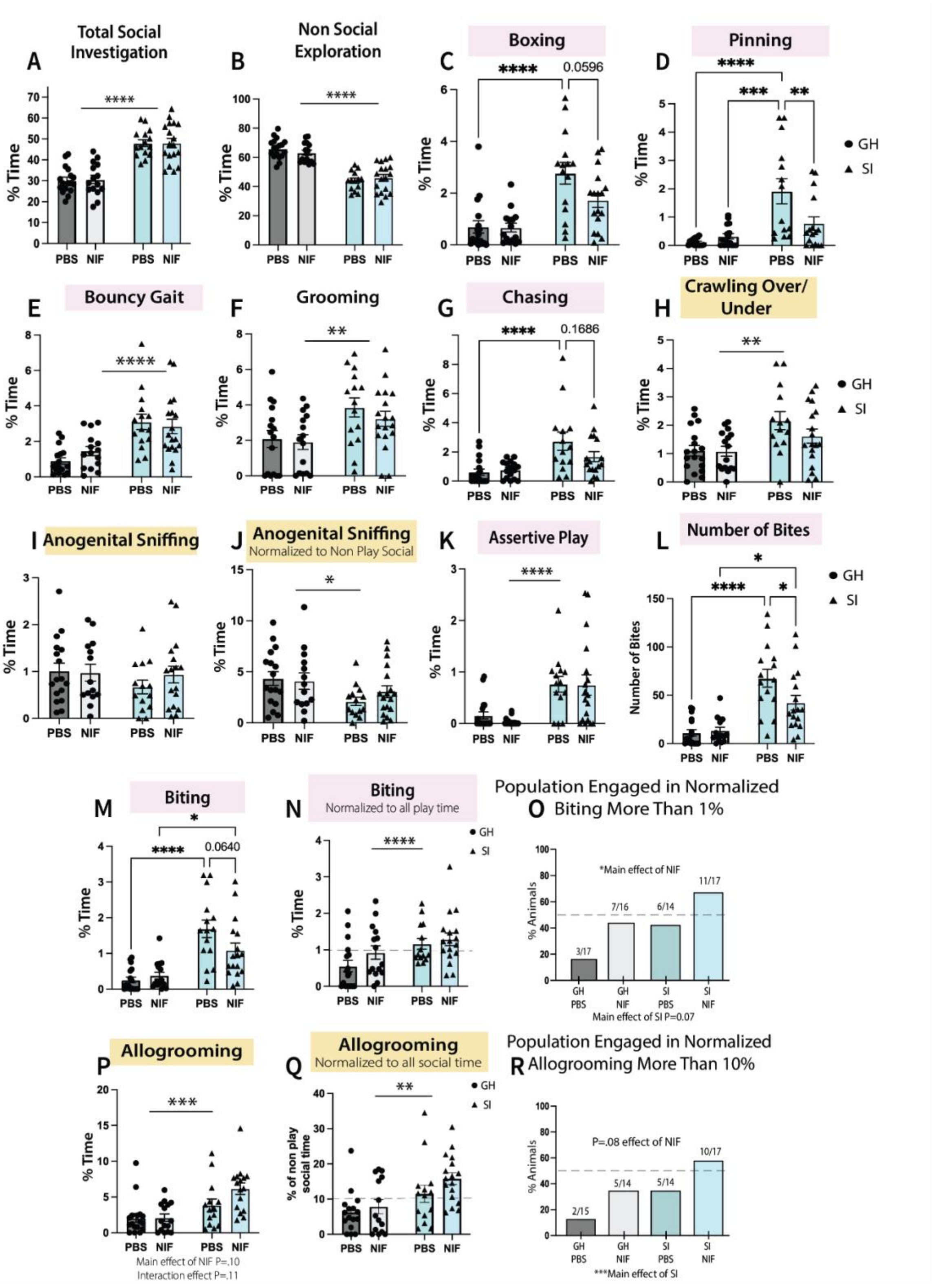


SF10: Individual Behavior Quantification Across NIF and SI. Quantification of behaviors represented as % time of total 15 minute play bout, dots represent a single animal, N=14-19 across at least 4 litters. A total social, B non-social exploration C boxing D pinning E bouncy gait

F grooming G chasing H crawling over/under I anogenital sniffing J anogenital sniffing normalized to non-play social K assertive play L number of bites M biting N biting normalized to all play time O population engaged in normalized biting by more than 1% P allogrooming Q allogrooming normalized to all social time R population engaged in normalized allogrooming by more than 10%. Significance by two-way ANOVA with Tukey's post hoc test *P<.05,**<.01,***<.001,****<.0001. Error bars = SEM.
